# Wall teichoic acid is required for DNA-triggered innate immune receptor activation by *Staphylococcus aureus*

**DOI:** 10.1101/2025.09.25.678139

**Authors:** Jordan B Jastrab, Ashley Tseng, Daniel Fisch, Stephanie A Ragland, Jonathan C Kagan

## Abstract

Receptors that stimulate inflammation are commonly activated by ligands that are buried within microbial cells. The mechanisms that facilitate immunostimulatory ligand release from microbes during infection are largely undefined. Using the pathogen *Staphylococcus aureus*, we describe wall teichoic acid (WTA) as a critical mediator of DNA release from bacterial cells during infection. Mechanistically, we found that the α-glycosylated form of WTA, generated by the enzyme TarM, is the primary mediator of bacterial DNA release within macrophages. This process is antagonized by the enzyme O-acetyltransferase, which prevents WTA attachment to peptidoglycan. This competition among peptidoglycan modifying enzymes determines activation of cytosolic DNA receptors AIM2 and cGAS. WTA therefore operates as a meta-regulator of inflammation, by controlling the availability of microbial ligands that stimulate innate immunity.

## INTRODUCTION

Central to immunity and host defense are pathogen-associated molecular patterns (PAMPs), which are microbial molecules with intrinsic immunostimulatory activities^1^. PAMPs have been under investigation for decades and are recognized as ligands for diverse families of innate immune pattern recognition receptors (PRRs)^2^. Among these PRRs are proteins that seed the assembly of inflammasomes, cytoplasmic protein complexes that regulate interleukin-1 (IL-1)-dependent inflammation^3^. Often in concert with Toll-like Receptor (TLR) signaling events, inflammasomes play key roles in resistance to infection and long-term immunity^4^.

Bacterial infections have informed much of our understanding of inflammasomes, with several PAMPs present in Gram-negative bacteria being identified as inflammasome-stimulatory factors (*e.g.* lipopolysaccharides (LPS)^5,6^, flagellin subunits^7,8^, and type 3 secretion system components^9^). While these factors bind directly to PRRs, the mechanisms by which these and other PAMPs are released from bacterial cells to access PRRs are largely undefined. This lack of knowledge is particularly evident when considering Gram-positive bacterial infections^10^, such as those caused by *Staphylococcus aureus* (*S. aureus*).

With the exception of *Mycobacterium tuberculosis*, *S. aureus* is thought to be the world’s leading bacterial cause of death^11^. Acute *S. aureus* infections are commonly associated with the production of bacterial pore-forming toxins (PFTs) such as α-hemolysin and LukAB^12^. The actions of these PFTs activate the inflammasome stimulatory protein NLRP3^13–15^, leading to interleukin-1β (IL-1β)-dependent inflammation^16–18^. In contrast to acute infections, persistent or chronic infections are associated with reduced PFT production through inactivation of the bacterial *agr* quorum sensing system^19,20^. Consequently, *agr*-inactive bacteria adopt a nontoxigenic phenotype^21^. Yet despite producing little to no PFTs, *agr*-inactive *S. aureus* are reported to activate inflammasomes^22,23^. Notably, human infections caused by nontoxigenic *agr*-inactive bacteria are associated with increased mortality^24^ and bacterial persistence^25–27^. The mechanisms by which nontoxigenic *S. aureus* influence inflammasomes and other innate immune signal transduction pathways are largely undefined.

Modification of peptidoglycan (PGN) by O-acetylation has emerged as one mechanism by which nontoxigenic *S. aureus* may avoid triggering inflammasomes. PGN is comprised of a polymer of repeating N-acetylmuramic acid (NAM) and N-acetylglucosamine (NAG) residues cross-linked by peptide bridges^28^. Many bacteria, including *S. aureus*, encode a PGN O-acetyltransferase (OatA) that catalyzes acetylation of the NAM C6 hydroxyl group^29^. O-acetylation increases PGN resistance to degradation by lysozyme^30,31^, an enzyme active within macrophage phagosomes that lyses bacteria by hydrolyzing the glycan linkage between NAM and NAG^32^. Whereas the role of OatA in lysozyme resistance *in vitro* is well documented^30,31^, no consensus immunological consequence of OatA activities during infection exists. For example, during infection of macrophages, *S. aureus oatA* mutants have been reported to display increased activation of inflammasomes, increased necroptosis, and increased likelihood of intraphagosomal bacteriolysis^22,33^. Additionally, NLRP3^14,22,23^, NLRP1^34^, NLRP6^35^, NLRP7^36^, and AIM2^33,37^ have each been independently reported to assemble inflammasomes during *S. aureus* infection or upon treatment with *S. aureus*-derived cell wall components. It therefore remains unclear how PGN O-acetylation impacts inflammasome activation, particularly in the context of infection with nontoxigenic bacteria.

In this study, we report that PGN modification by wall teichoic acid (WTA) is required for inflammasome activation induced by multiple laboratory and clinical strains of *S. aureus*. This process is enhanced by WTA glycosylation via the bacterial enzyme TarM. PGN O-acetylation by OatA reduces WTA levels in *S. aureus*, thus explaining the inflammasome-inhibitory activities of OatA. Mechanistically, TarM and OatA differentially regulate the amount of bacterial extracellular DNA released into infected macrophages after phagocytosis, thus controlling activation of the cytosolic DNA sensors AIM2 and cGAS. WTA is therefore not an intrinsically inflammatory entity, akin to a PAMP, but is rather an example of a distinct class of bacterial factors that promote PAMP release and innate immune receptor activation.

## RESULTS

### Lysozyme does not regulate IL-1β release during *S. aureus* infection of macrophages

To explore how nontoxigenic *S. aureus* influences inflammasome activities, studies were performed with *S. aureus* strain SA113, in which the *agr* system is permanently inactive^38^. To evaluate the role of OatA, we obtained a previously described SA113 *oatA* mutant containing an Δ*oatA*::*kan* deletion/insertion mutation^22,31^. Unexpectedly, we found that this SA113 *oatA* strain was toxigenic, as it displayed hemolytic activity on blood agar plates, and exhibited a slight growth advantage over its parental wild-type (WT) strain (Fig S1A,B). Neither phenotype was complemented by reintroduction of the *oatA* gene (Fig S1A,B). These findings suggested that this mutant strain was not an ideal model to study nontoxigenic activities of *S. aureus*. We therefore rederived the *oatA* mutant (SA_Δ_*_oatA_*) by transducing the Δ*oatA*::*kan* mutation into WT SA113 (SA_WT_). Our rederived SA_Δ_*_oatA_* exhibited similar growth in tryptic soy broth (TSB) as SA_WT_ (Fig S1C) and was non-hemolytic on blood agar (Fig S1D). While we reproduced findings that the previously generated *oatA* mutant exhibits a large decrease in intracellular survival within macrophages^22^, the rederived SA_Δ_*_oatA_* exhibited only a small survival defect at late infection timepoints (Fig S1E). To avoid confounding OatA-independent phenotypes, we focused our studies on this newly derived SA_Δ_*_oatA_* mutant.

We infected primary bone marrow-derived macrophages (BMDMs) with SA_WT_ or SA_Δ_*_oatA_.* Extracellular medium from infected cells was then measured for release of IL-1β to assess inflammasome activation and LDH activity to measure pyroptotic cell death. As expected^22^, we found that SA_Δ_*_oatA_* triggered the release of more IL-1β and LDH than SA_WT_ (Fig 1A,B). In contrast, minimal strain-specific differences were detected when we measured TNFα secretion, a common readout for TLR signaling^39^ (Fig 1C). Priming of cells by pretreatment with LPS, a TLR4 ligand that induces pro-IL-1β production, increased the total amounts of IL-1β released but did not alter the trends observed between bacterial strains (Fig 1A).

**Figure 1.**
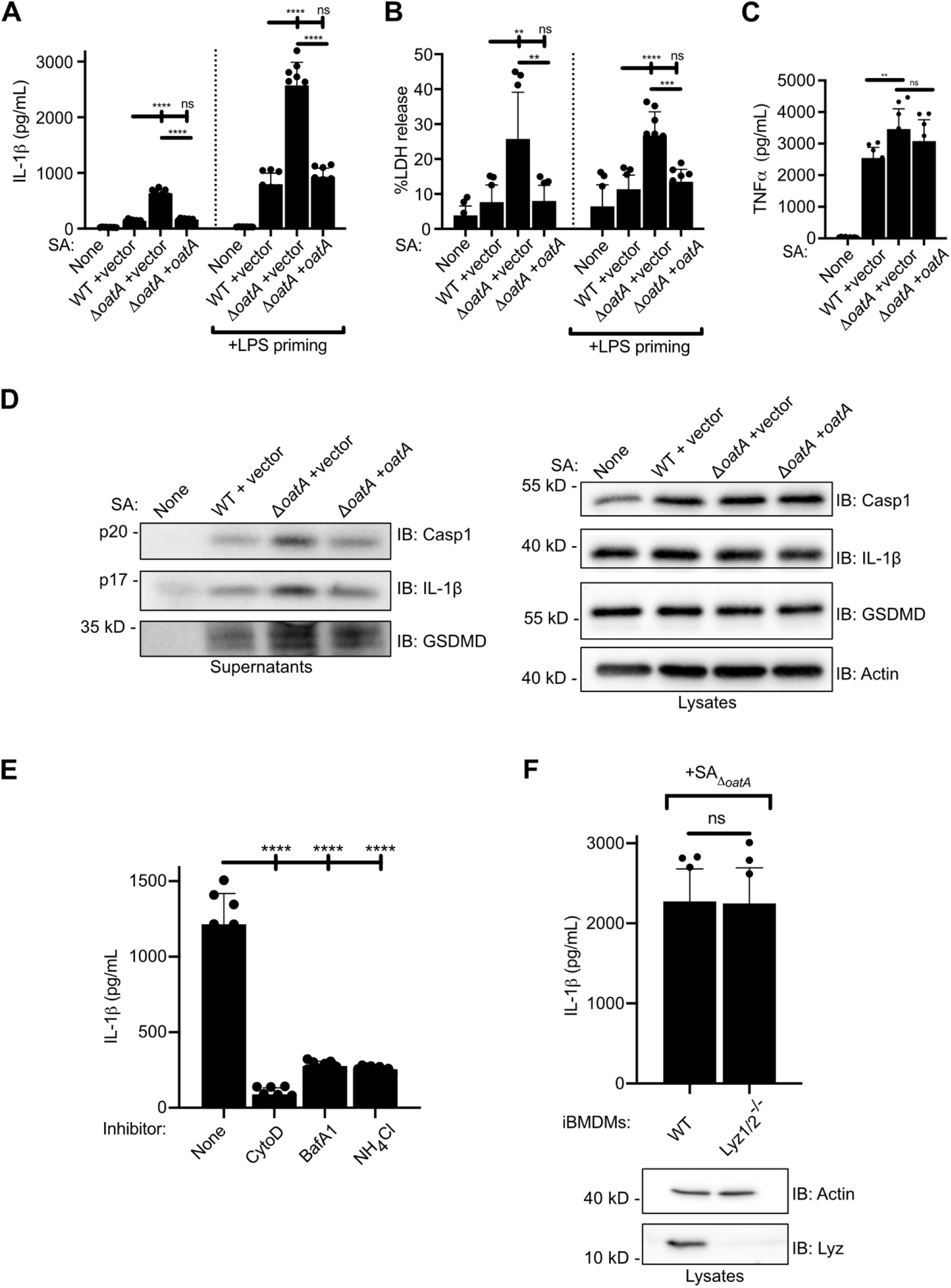
*S. aureus* OatA blunts lysozyme-independent inflammasome activation during nontoxigenic infection of macrophages. (A-C) WT BMDMs were primed where indicated with 1 µg/mL LPS for 3 hours. Overnight cultures of the indicated *S. aureus* strains were washed in sterile PBS and added to BMDMs at a multiplicity of infection (MOI) of 10. 30 minutes post-infection, supernatants were replaced with media containing 20 µg/mL lysostaphin to kill non-phagocytosed bacteria. 4 hours post-infection (hpi), supernatants were collected and analyzed by (A) Lumit assay for IL-1β, (B) LDH assay, or (C) ELISA for TNFα. (D) WT BMDMs were primed with LPS and infected as above with the indicated *S. aureus* strains at an MOI of 10. 4 hpi, supernatants and lysates were collected, separated by SDS-PAGE, and immunoblotted (IB) for the indicated proteins. p20 and p17 represent the cleaved activated forms of caspase-1 (Casp1) and IL-1β, respectively; the cleaved gasdermin D (GSDMD) N-terminal domain runs at approximately 32 kD. (E) WT iBMDMs were primed with LPS and infected with SA_Δ_*_oatA_* at an MOI of 10. The indicated inhibitors were added 30 minutes prior to infection and maintained for the remainder of the experiment. To maintain the presence of non-phagocytosed bacteria, lysostaphin was not added. 4 hpi, supernatants were collected and analyzed by Lumit assay for IL-1β. (F) The indicated iBMDM lines were primed with LPS and infected with SA_Δ_*_oatA_* as above. 4 hpi, supernatants were collected and analyzed by Lumit assay for IL-1β. For immunoblots, whole cell lysates of uninfected cells were separated by SDS-PAGE and immunoblotted for lysozyme (Lyz) or β-actin. kD: kilodaltons.

To complement these analyses, we performed immunoblot analysis of extracellular media from infected cells to assess the cleaved (*i.e.* active) forms of key inflammasome effectors caspase-1, IL-1β, and gasdermin D. Infections with SA_Δ_*_oatA_* yielded the highest amounts of these cleaved proteins (Fig 1D). A complemented strain in which a single chromosomal copy of the *oatA* gene was reintroduced under control of its native promoter (SA_Δ_*_oatA +oatA_*) phenocopied SA_WT_ in all assays (Fig 1A-D, Fig S1C-E).

OatA is best recognized as an enzyme that acetylates PGN, rendering the cell wall resistant to the activity of lysozyme^30^. Because lysozyme is active within acidified phagolysosomes^32^, it has been generally accepted that lysozyme sensitivity determines the inflammasome activities induced by *S. aureus oatA* mutants. However, the role of lysozyme during *S. aureus* infection has only been explored using chemical inhibitors^22,33^. To determine if *S. aureus* requires passage through the phagolysosomal system to activate the inflammasome, we chemically inhibited phagocytosis with cytochalasin D (CytoD) or phagolysosomal acidification with either bafilomycin A1 (BafA1) or ammonium chloride (NH_4_Cl). Each of these treatments blocked IL-1β production during infection of immortalized BMDMs (iBMDMs) (Fig 1E). To clarify the role of lysozyme in the activation of inflammasomes by *S. aureus*, we generated iBMDMs with disruptions in the genes encoding both isoforms of lysozyme (Lyz1/2^-/-^) using CRISPR-Cas9. The resulting macrophages contained no detectable lysozyme protein by immunoblotting (Fig 1E, bottom). Unexpectedly, lysozyme deficiency did not impact IL-1β release during SA_Δ_*_oatA_* infection (Fig 1E, top). These data indicate that *S. aureus* OatA prevents inflammasome activation via a mechanism that is distinct from its established role in blocking lysozyme activity.

### A genome-wide genetic screen in *S. aureus* identifies wall teichoic acid as an inflammasome regulatory factor

Having eliminated the lysozyme pathway as a host factor that regulates inflammasomes during *S. aureus* infection, we sought to identify new mechanisms that mediate this process. Towards this end, we performed a forward genetic screen to identify bacterial mutants that are defective for triggering inflammasome activities (Fig S2A). We used the mariner-based *bursa aurealis* transposon^40^ to generate random insertions at TA sites across the bacterial genome. As all well-defined inflammasome assays require population-based measurements (*e.g.* ELISA or immunoblotting), screening of the transposon library required an infection format enabling single bacterial strain assessments. We devised a system whereby iBMDMs were seeded into 384-well plates, and each well was infected with an individual bacterial mutant. IL-1β production from each well was assessed using the Lumit chemiluminescent assay^41^, which can be performed *in situ* (Fig S2A). To maximize the dynamic range of IL-1β production detected in this screen, the SA_Δ_*_oatA_* mutant was selected as the parent strain for transposon mutagenesis. Using this system, we identified transposon mutant 3H4, which was defective for IL-1β and LDH release as compared to the parental SA_Δ_*_oatA_* strain (Fig 2A-B). Whole genome sequencing of this strain revealed a single transposon insertion in the operon encoding *tarM* (Fig S2B), a gene not previously described to have a role in inflammasome function. Complementation of 3H4 with *tarM* restored IL-1β and LDH release to levels comparable to the parental SA_Δ_*_oatA_* strain (Fig 2A,B). In contrast, disrupting *tarM* did not impact infection-induced TNFα secretion (Fig 2C).

**Figure 2.**
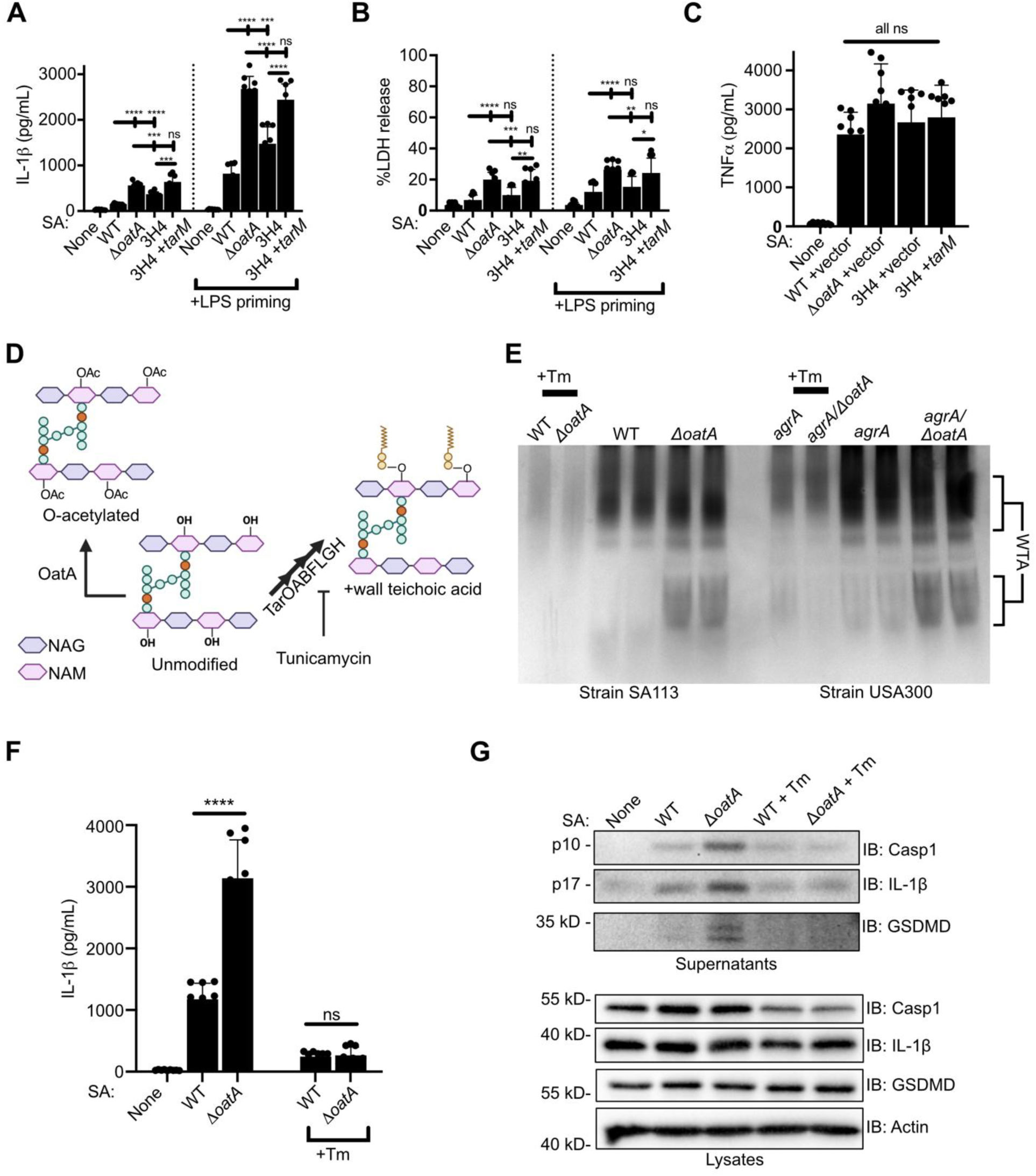
A forward genetic screen reveals that wall teichoic acid is required for activation of inflammasomes during nontoxigenic *S. aureus* infection. (A-C) WT BMDMs were primed where indicated with LPS and then infected with the indicated *S. aureus* strains at an MOI of 10 as described above. 4 hpi, supernatants were collected and analyzed by (A) Lumit assay for IL-1β, (B) LDH assay, or (C) ELISA for TNFα. (D) Cartoon depicting the location of PGN at which O-acetylation and WTA attachment occurs. TarOABFLGH: enzymes responsible for synthesis and deposition of WTA. (E) The indicated *S. aureus* strains were grown to stationary phase and sacculus was prepared. Where indicated, bacteria were grown in the presence of 0.4µg/mL tunicamycin to inhibit WTA synthesis. WTA was hydrolyzed from purified sacculus by treatment with sodium hydroxide, separated by non-reducing and non-denaturing PAGE, and visualized by silver staining. (F,G) WT BMDMs were primed with LPS and infected with the indicated *S. aureus* strains at an MOI of 10 as described above; *S. aureus* was grown with or without 0.4 µg/mL tunicamycin as indicated. 4 hpi, supernatants and whole cell lysates were collected. Supernatants were analyzed by Lumit assay for IL-1β (F). Supernatants and lysates were separated by SDS-PAGE and analyzed by immunoblotting (H).

TarM is a glycosyltransferase that modifies wall teichoic acid (WTA)^42^, a PGN-anchored polymer that is crucial for regulating bacterial growth and morphology^43^. Notably, the attachment site for WTA onto PGN is the identical site that is acetylated by OatA^43^ (Fig 2D). The observation that a shared PGN site can be modified either by WTA or acetylation suggested an enzymatic competition for this attachment site, the outcome of which could impact inflammasome activities. A prediction of this model is that WTA should be differentially abundant on bacterial strains that contain or lack OatA. To test this prediction, we purified sacculus from SA_WT_ and SA_Δ_*_oatA_*, released WTA from PGN by treatment with sodium hydroxide, and visualized solubilized WTA by polyacrylamide gel electrophoresis (PAGE) and silver staining. As a control, we grew bacteria in the presence of tunicamycin (Tm), an inhibitor of WTA synthesis^44^. Consistent with our hypothesis, SA_Δ_*_oatA_* contained higher amounts of WTA than SA_WT_ (Fig 2E). Similar results were obtained using a distinct set of *S. aureus* strains derived from the methicillin-resistant *S. aureus* (MRSA) USA300 JE2 background (Fig 2E). While SA_Δ_*_oatA_* contained higher amounts of WTA-modified PGN, it did not exhibit an increase in expression of the WTA synthetic or glycosyltransferase genes *tarO*, *tarM*, or *tarS* (Fig S2C).

Based on the relationship between OatA and WTA abundance, we examined the influence of WTA on inflammasome activities during infection. BMDMs were infected with bacteria grown in the presence or absence of tunicamycin to block WTA synthesis. Depletion of WTA by tunicamycin ablated IL-1β release overall and eliminated the difference in secretion of IL-1β elicited by SA_WT_ and SA_Δ_*_oatA_* (Fig 2E). WTA had a similar effect on the production of cleaved IL-1β, caspase-1, and gasdermin D during infection as evaluated by immunoblotting (Fig 2F).

### WTA glycosylation maximizes inflammasome activation during *S. aureus* infection

The above findings prompted mechanistic investigations of the link between inflammasomes and the WTA glycosyltransferase TarM. The generation of glycosylated WTA (glycoWTA) in *S. aureus* occurs by the addition of NAG to WTA by at least three distinct glycosyltransferases: TarS, TarM, and TarP (Fig 3A). The *tarS* gene is present in nearly all *S. aureus* strains^45^ and is encoded within an operon that includes core WTA synthetic genes. In contrast, TarM and TarP are believed to be accessory glycosyltransferases whose presence varies across *S. aureus* clonal complexes^45^. We confirmed by whole genome sequencing that strain SA113 encodes *tarS* and *tarM* but not *tarP* (not shown). TarS, TarM, and TarP attach NAG to WTA via β(1,4)^46^, α(1,4)^42^, or β(1,3)^47^ linkages, respectively. Like TarM, the functions of these other WTA glycosyltransferases in the regulation of inflammasome activity are unknown.

**Figure 3.**
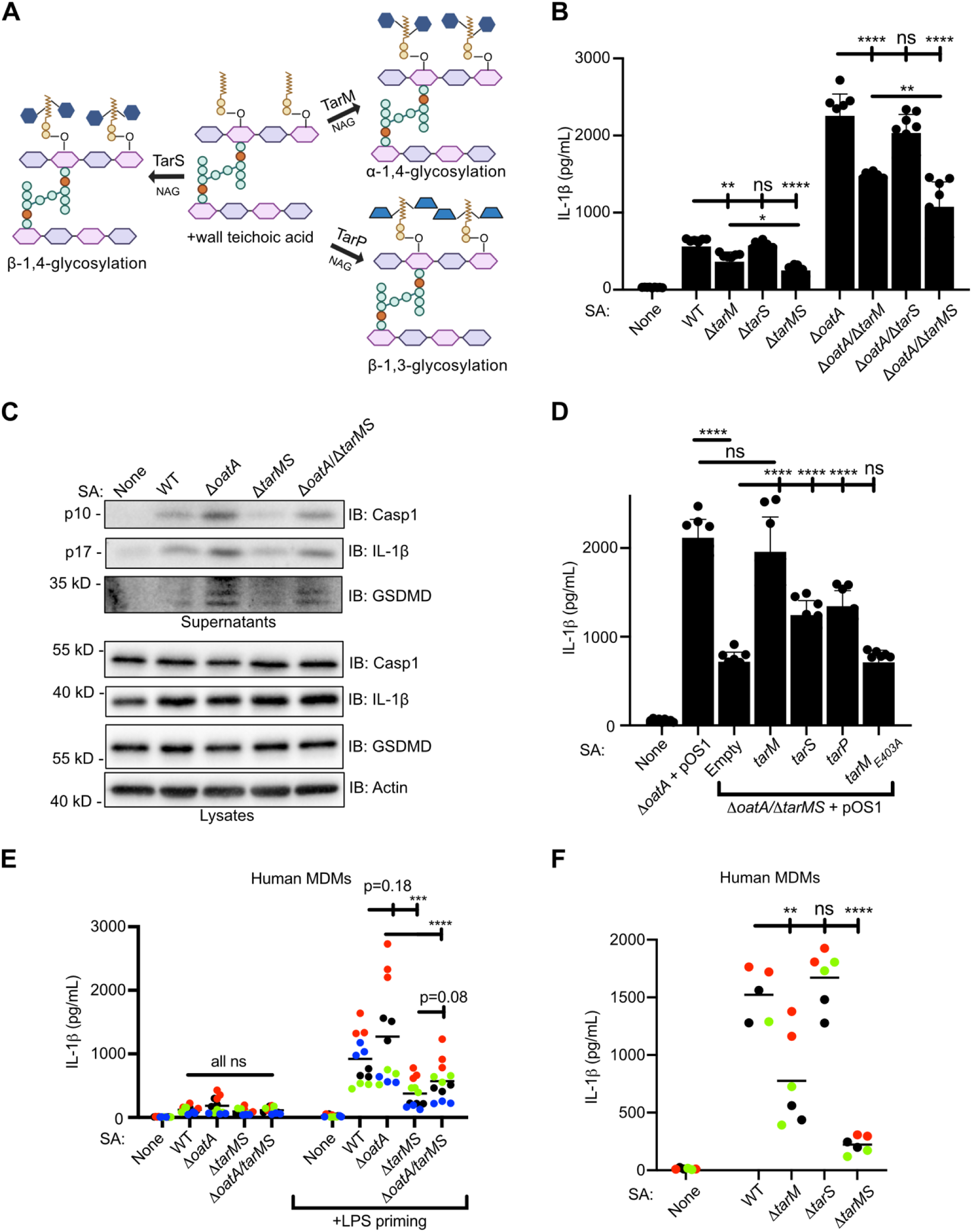
*S. aureus* WTA glycosyltransferases promote inflammasome activation. (A) Cartoon depicting the activities of known *S. aureus* WTA glycosyltransferases. (B,C) WT BMDMs were primed with LPS and infected with the indicated *S. aureus* strains at an MOI of 10 as described above. 4 hpi, supernatants and whole cell lysates were collected. Supernatants were analyzed by Lumit assay for IL-1β (B). Supernatants and lysates were separated by SDS-PAGE and analyzed by immunoblotting (C). (D) WT BMDMs were primed with LPS and infected with the indicated *S. aureus* strains at an MOI of 10 as described above. 4 hpi, supernatants were collected and analyzed by Lumit assay for IL-1β. (E,F) hMDMs were primed where indicated with LPS and then infected with the indicated *S. aureus* strains at an MOI of 10 as described above. 4 hpi, supernatants were collected and analyzed by ELISA for IL-1β. Similarly colored dots within each experiment represent technical replicates using cells from the same donor.

To determine the relationship between TarM and TarS activities on the activation of inflammasomes, we generated unmarked, in-frame deletions of *tarM*, *tarS*, or both in the SA_WT_ and SA_Δ_*_oatA_* backgrounds. All bacterial strains grew comparatively *in vitro* (Fig S2D). BMDMs infected with a Δ*tarM* mutant (SA_Δ_*_tarM_*) displayed reduced IL-1β production compared to SA_WT_, whereas deletion of *tarS* to generate strain SA_Δ_*_tarS_* did not impact IL-1β production significantly. Notably, BMDMs infected with a strain lacking both *tarM* and *tarS* (SA_Δ_*_tarMS_*) produced the least amount of IL-1β. Examination of Δ*tarM* and Δ*tarS* mutants in the SA_Δ_*_oatA_* background demonstrated similar Δ*tarM-*, Δ*tarS-,* and Δ*tarMS*-specific phenotypes (Fig 3B). To complement the above analyses, we evaluated supernatants of infected cells by immunoblotting, which demonstrated that strains lacking *tarMS* triggered less release of active caspase-1, gasdermin D, and IL-1β than *tarMS-*sufficient strains (Fig 3C). In contrast to these inflammasome-associated phenotypes, WTA glycosyltransferase gene deletions had minimal impact on TNFα secretion (Fig S2E). *S. aureus* WTA can also be covalently modified by alanylation. Although on multiple attempts we could not obtain deletion mutants in the *dltABCD* operon responsible for WTA alanylation^48^, deletion of *graRS*, which encodes the two-component system that regulates *dltABCD* expression^49^, had no effect on IL-1β release during infection (Fig S2F).

To complement the deficiencies of each glycosyltransferase, we transformed the SA_Δ_*_oatA_*_/Δ_*_tarMS_* mutant with plasmids encoding individual glycosyltransferase genes under control of the constitutive P*_lgt_* promoter^50^. Expression of *tarM* fully rescued IL-1β release to the level of SA_Δ_*_oatA_*, whereas expression of *tarS* or *tarP* under the same promoter provided only partial rescue (Fig 3D). Expression of the *tarM_E403A_* allele, which includes a point mutation that renders TarM catalytically inactive^51^, did not rescue IL-1β release.

To determine if glycoWTA impacts inflammasome activation in human cells, we infected primary human monocyte-derived macrophages (hMDMs). Consistent with our findings in mouse macrophages, Δ*tarMS* mutants triggered less IL-1β production as compared to *tarMS*-sufficient counterparts (Fig 3E) even though Δ*tarMS* strains triggered slightly more TNFα secretion (Fig S2F). Although Δ*oatA* strains demonstrated a trend towards driving more IL-1β release than *oatA*-sufficient strains (Fig 3E), this effect was not statistically significant and deletion of *oatA* did not impact TNFα secretion (Fig S2G). Finally, we infected hMDMs with single gene deletion strains and found that deletion of *tarS* alone had no impact on IL-1β release whereas a Δ*tarM* mutant triggered reduced IL-1β release as compared to WT bacteria (Fig 3F). Δ*tarMS* mutants triggered the least IL-1β release as compared to all other strains examined (Fig 3F). These data are consistent with findings in BMDMs and suggest that WTA glycosylation enhances activation of inflammasomes during nontoxigenic *S. aureus* infection, with α(1,4)-glycoWTA playing a dominant role in promoting inflammasome activity.

### AIM2 mediates inflammasome activation during nontoxigenic *S. aureus* infection

Although at least five inflammasomes have been reported to promote IL-1β production during *S. aureus* infection^33,35,36,52,53^, the NLRP3 inflammasome has been implicated most frequently. However, work in this area has focused on activities in response to bacterial PFTs^14–16,18,54,55^ or upon treatment with purified PGN^22,23,56^. To examine this question during infections, we isolated BMDMs from mice deficient in NLRP3, AIM2, or the inflammasome-associated IL-1β converting enzyme caspase-1. Treatments of BMDMs with the NLRP3 agonist nigericin^52^ or the AIM2 agonist poly(dA:dT)^57^ confirmed the expected NLRP3-dependent and AIM2-dependent activities, respectively (Fig S3A).

Infections of these genotypes of BMDMs with SA_Δ_*_oatA_* demonstrated that caspase-1 was required for IL-1β and LDH release (Fig 4A) and for production of cleaved gasdermin D and IL-1β (Fig 4B). In contrast, NLRP3 was not required for the activation of these inflammasome activities nor for production of cleaved caspase-1 (Fig 4A,B). Similarly, chemical inhibition of NLRP3 with MCC950^58^ blocked IL-1β release during treatment with nigericin but had no effect during SA_Δ_*_oatA_* infection (Fig S3B). Notably, AIM2 was required for IL-1β release and cleavage of caspase-1, gasdermin D, and IL-1β in response to SA_Δ_*_oatA_* infection (Fig 4A,B). These data indicate that whereas NLRP3 is the primary mediator of inflammasome activities induced by PFTs and purified peptidoglycan^13,23^, AIM2 is the primary mediator of inflammasome activation during nontoxigenic *S. aureus* infection.

**Figure 4.**
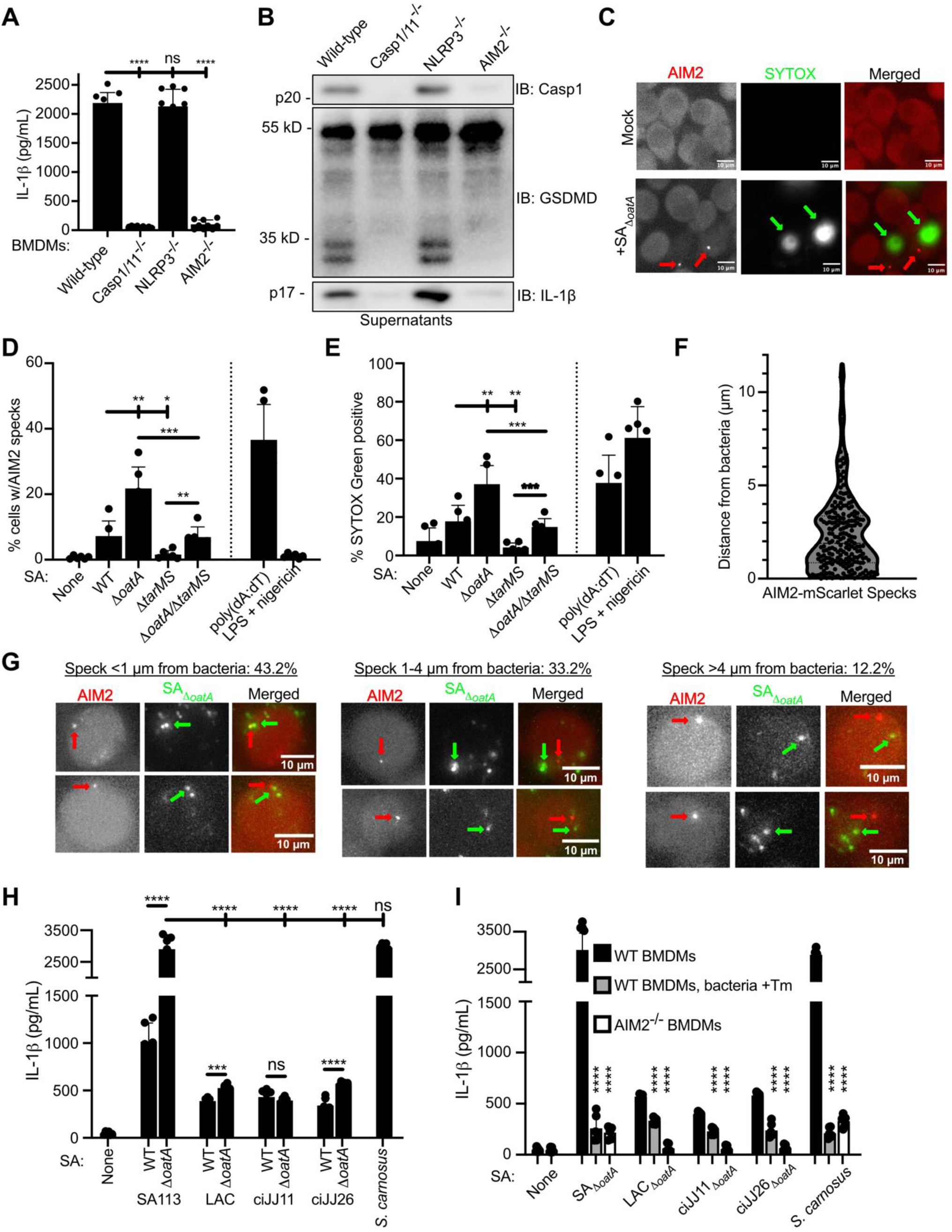
Non-toxigenic *S. aureus* activates the AIM2 inflammasome in BMDMs. (A) BMDMs generated from the indicated mouse lines were primed with LPS and infected with SA_Δ_*_oatA_* at an MOI of 10 as described above. 4 hpi, supernatants were collected and analyzed by Lumit assay for IL-1β. (B) BMDMs generated from the indicated mouse lines were primed with LPS and infected with SA_Δ_*_oatA_* at an MOI of 10 as described above. 4 hpi, supernatants and whole cell lysates were collected, separated by SDS-PAGE, and analyzed by immunoblotting. (C-E) Unprimed AIM2^-/-^-AIM2-mScarlet iBMDMs were infected with the indicated *S. aureus* strains at an MOI of 10; 30 minutes post-infection lysostaphin and SYTOX green were added to the infection mixture. Infections were imaged at 1-minute intervals by fluorescence widefield microscopy for a total of 4 hours. (C) Representative images of mock infected (top) or SA_Δ_*_oatA_* - infected (bottom) cells; red arrows indicate AIM2 specks, green arrows indicate SYTOX Green-positive nuclei. (D,E) Quantification of the percentage of total cells that developed AIM2 specks (D) or SYTOX Green uptake (E) over 4 hours of infection. (F,G) Unprimed AIM2^-/-^-AIM2-mScarlet iBMDMs were pretreated with zVAD-FMK and infected with SA_Δ_*_oatA_* at an MOI of 10. 30 minutes post-infection, lysostaphin was added to infection media. 5 Z-stack images were obtained at 1-minute intervals by fluorescence widefield microscopy for approximately 3 hours; stacks were projected into a single image and the distance to the nearest bacterium was measured for each nascent AIM2 speck. (F) Box and violin plot noting the distance between each AIM2 speck and the nearest detectable bacterium. n=460 puncta from 3 independent experiments. (G) Representative images of AIM2 specks that formed <1 µm (left), 1-4 µm (middle), or >4 µm (right) from a bacterium. Red arrows: AIM2 specks; Green arrows: closest detectable bacterium. (H) WT BMDMs were primed with LPS and infected with the indicated staphylococcal strains at an MOI of 10 as described above. 4 hpi, supernatants were collected and analyzed by Lumit assay for IL-1β. (I) WT or AIM2^-/-^ BMDMs were primed with LPS and infected with the indicated staphylococcal strains at an MOI of 10 as described above; where indicated, bacteria were grown in the presence of 0.4µg/mL (SA113) or 0.8µg/mL (other strains) tunicamycin. 4 hpi, supernatants were collected and analyzed by Lumit assay for IL-1β.

To characterize inflammasome activation by *S. aureus* at the single cell level, we generated AIM2^-/-^ iBMDMs and rescued AIM2 activity by transducing cells with a gene encoding an AIM2-mScarlet fluorescent fusion protein (Fig S3C). Infection of AIM2-mScarlet iBMDMs with SA_Δ_*_oatA_* or treatment with poly(dA:dT) led to the formation of dense AIM2-mScarlet specks within cells (Fig 4C,D), akin to the behavior of the inflammasome adaptor ASC in other studies^59,60^. AIM2-mScarlet specks were not detected in cells treated with LPS + nigericin (Fig 4D). Consistent with the bacterial genotype-specific trends observed in our population-based studies, we found that infections with SA_WT_, SA_Δ_*_oatA_*, SA_Δ_*_tarMS_*, and SA_Δ_*_oatA/_*_Δ_*_tarMS_* led to the formation of AIM2 specks in 7.2%, 21.7%, 1.5%, or 6.9% of cells, respectively (Fig 4D). A similar pattern was observed for cell death, as indicated by uptake of the cell-impermeable nucleic acid stain SYTOX green (Fig 4E).

To define the spatial dynamics of AIM2 inflammasome activation, AIM2-mScarlet iBMDMs were infected with a strain of SA_Δ_*_oatA_* expressing green fluorescent protein (GFP). Infected cells were treated with the pan-caspase inhibitor zVAD-FMK to ensure nascent AIM2 inflammasomes were not lost upon pyroptosis, as described previously^61^. We found that 43.2% of AIM2 specks formed within 1 µm of a bacterium, 33.2% formed within 1-4 µm, 12.2% formed >4 µm from a bacterium, and 11.4% formed in cells with no detectable bacteria (Fig 4F,G). Thus, nearly 90% of AIM2 inflammasomes formed within detectably infected cells, with the largest proportion forming in close proximity to a bacterial cell.

### Virulent clinical strains of *S. aureus* activate AIM2

To determine the role of AIM2 during infection with clinically relevant bacteria, we expanded our studies to include ciJJ11, a clinical MRSA isolate obtained from the blood of a patient with relapsed prosthetic valve endocarditis, and ciJJ26, a clinical MRSA isolate obtained from the blood of a patient with extensive extracellular graft material who developed recurrent bacteremia. Additionally, we examined LAC^62^, a well-studied USA300 MRSA isolate. Each of these virulent bacterial strains was nontoxigenic under our infection conditions, as infection of BMDMs elicited minimal cytotoxicity (Fig S3D). To determine if these clinical isolates utilize PGN O-acetylation to reduce inflammasome activity, we generated in-frame, unmarked Δ*oatA* deletions in each virulent isolate. Compared to their WT parental strains, LAC_Δ_*_oatA_* and ciJJ26_Δ_*_oatA_* triggered increased IL-1β release, whereas deletion of *oatA* from ciJJ11 had no effect (Fig 4H). In contrast to the variable impact of OatA, tunicamycin treatment to deplete WTA reduced IL-1β production during infection with all bacterial strains examined (Fig 4I). Additionally, AIM2 was required for the release of IL-1β in response to infection with all examined strains (Fig 4I).

We noted that all virulent clinical strains of *S. aureus* triggered substantially less IL-1β production than the attenuated SA113-derived strains. To determine if this trend towards hyper-engagement of AIM2 by non-virulent bacteria extends beyond *S. aureus*, we infected BMDMs with *Staphylococcus carnosus* (*S. carnosus*), a non-virulent staphylococcal species that is naturally deficient in PGN O-acetylation^63^. Similar to SA_Δ_*_oatA_*, *S. carnosus* triggered high amounts of IL-1β release (Fig 4H) in a manner dependent on WTA and AIM2 (Fig 4I). All bacterial strains induced comparable amounts of TNFα secretion (Fig S3E), suggesting that the differences in IL-1β production between virulent and non-virulent strains are not explained by differential TLR signal transduction. Thus, regardless of the magnitude of inflammasome-stimulatory activities, WTA and AIM2 are common mediators of inflammasome activation by virulent clinical strains and non-virulent strains of *Staphylococci*.

### Cell wall modifications control the availability of extrabacterial DNA for multiple pattern recognition receptors

To define the mechanism by which OatA and TarMS regulate AIM2 activity, we determined if these enzymes might alter internalization of *S. aureus* by BMDMs. However, there was no significant difference in the percentage of BMDMs containing internalized bacteria when comparing infection with SA_WT_ to infection with SA_Δ_*_oatA_* or SA_Δ_*_tarMS_* (Fig 5A). Another explanation for the different inflammasome phenotypes between bacterial strains is differential bacterial killing after phagocytosis. However, quantifying intracellular survival of bacteria during infection of BMDMs revealed no difference between SA_WT_, SA_Δ_*_oatA_*, and SA_Δ_*_tarMS_* (Fig 5B).

**Figure 5.**
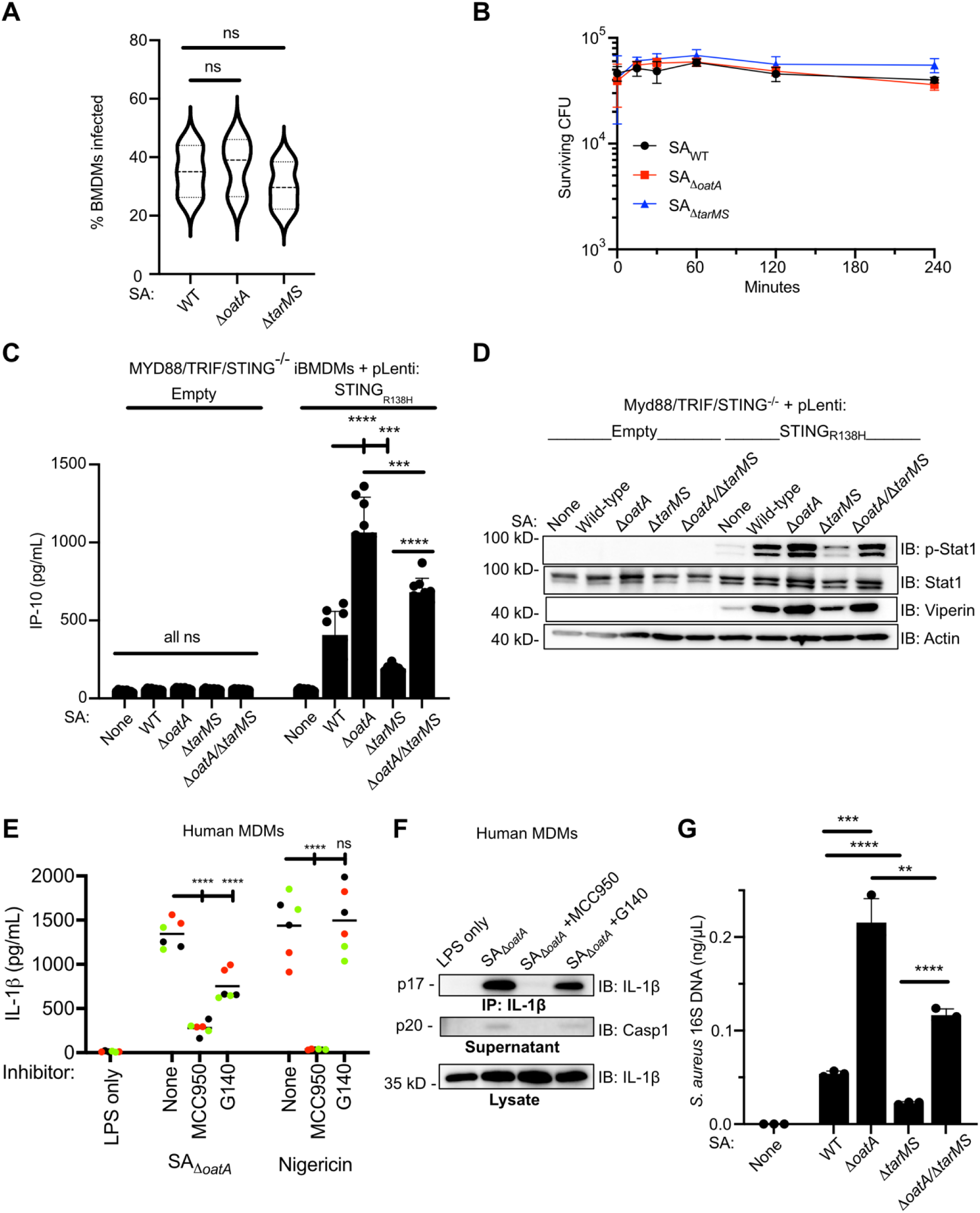
*S. aureus* OatA and TarMS alter extrabacterial DNA availability within macrophages to regulate the activities of multiple cytosolic DNA sensors. (A) Unprimed WT BMDMs were plated on a glass coverslip and infected with the indicated *S. aureus* strains at an MOI of 1. 1 hpi, cells were fixed with 4% PFA and washed extensively with PBS. Extracellular S*. aureus* was stained green in the absence of a permeabilizing agent, cells were fixed again, and total *S. aureus* was subsequently stained red in the presence of a permeabilizing agent. Stained cells were imaged by confocal microscopy and the percentage of cells containing intracellular bacteria (stained red but not green) was quantified. (B) Unprimed WT BMDMs were infected with the indicated *S. aureus* strains at an MOI of 1; where indicated, 30 minutes post-infection lysostaphin was added to a final concentration of 20 µg/mL. At the indicated time points, cells were washed with PBS and lysed with 1% saponin to release intracellular bacteria; bacteria were serially diluted and plated on solid media to enumerate surviving CFUs. Data are representative of 3 independent experiments. (C) Unprimed iBMDM lines were infected with the indicated *S. aureus* strains at an MOI of 10 as described above. 4 hpi, supernatants were collected and analyzed by ELISA for IP-10. (D) Unprimed iBMDMs were infected with the indicated *S. aureus* strains at an MOI of 10 as described above. 4 hpi, whole cell lysates were prepared, separated by SDS-PAGE, and analyzed by immunoblotting. (E) hMDMs were primed with LPS and either infected with SA_Δ_*_oatA_* at an MOI of 10 for 4 hours or treated with 10 µM nigericin for 2 hours. Where indicated, cells were pretreated for 30 min and subsequently maintained in 5 µM MCC950 or 20 µM G140. Supernatants were collected and analyzed by ELISA for IL-1β. (F) hMDMs were primed with LPS and infected with SA_Δ_*_oatA_* at an MOI of 10 for 4 hours in the presence or absence of MCC950 or G140. 4 hpi, supernatants and whole cell lysates were prepared, an aliquot of supernatant was removed, and the remainder was subjected to anti-IL-1β immunoprecipitation (IP). Supernatants, IP eluates, and lysates were separated by SDS-PAGE and analyzed by immunoblotting. (G) AIM2^-/-^ iBMDMs were infected with the indicated *S. aureus* strains at an MOI of 50, and after 30 minutes infection media was replaced with media containing 20 µg/mL lysostaphin. 3 hours post-infection, cells were washed with PBS, lysed with radioimmunoprecipitation assay (RIPA) buffer, and intact bacteria were removed by centrifugation. DNA was purified from cell lysates and bacterial DNA was quantified by qPCR using primers specific to the gene encoding *S. aureus* 16S RNA. DNA levels were normalized to total host cell DNA as measured by qPCR using primers specific to the gene encoding mouse β-actin. Data are representative of 3 independent experiments.

The finding that OatA and TarMS do not impact bacterial internalization or survival suggested these factors may instead alter the release of other immunogenic molecules from intracellular bacteria. As AIM2 is a cytosolic dsDNA sensor, we hypothesized that these cell wall modifying enzymes alter the release of DNA into the host cell cytosol. If OatA and TarMS impact DNA release into the cytosol, these enzymes should impact the activities of additional cytosolic DNA sensors, such as cyclic GMP-AMP synthase (cGAS). cGAS is the principal innate immune receptor that drives interferon (IFN) responses to cytosolic DNA^64^. Upon binding DNA, cGAS converts into an active enzyme that produces the cyclic dinucleotide 2’,3’-cyclic GMP-AMP (cGAMP), which activates the protein Stimulator of Interferon Genes (STING) to trigger downstream IFN responses^65^. To address this possibility, we infected iBMDMs that were engineered such that cGAS-STING is the only pathway capable of activating IFN gene expression. Towards this end, MyD88, TRIF, STING triple knockout iBMDMs (MyD88/TRIF/STING^-/-^) were rescued with a transgene encoding STING_R138H_, a variant that is activated by cGAS-derived cGAMP but not by bacteria-derived cyclic dinucleotides, including those produced by *S. aureus*^66,67^. These cells were then infected with *S. aureus* strains. Consistent with our observations of AIM2 activities, infections with SA_Δ_*_oatA_* triggered the highest activation of cGAS-STING as measured by secretion of the IFN-stimulated factor IP-10, while Δ*tarMS* strains triggered the lowest amount of IP-10 release (Fig 5C). Similar results were obtained by immunoblot analysis for the phosphorylation of IFN-responsive transcription factor STAT-1 and production of the IFN-stimulated factor viperin (Fig 5D). In the absence of STING_R138H_, no IFN activities were evident during any infection, thus confirming the fidelity of this genetic system as a reporter on cGAS-STING function. Two distinct cytosolic DNA sensors, AIM2 and cGAS, are therefore similarly altered by OatA and TarMS.

In primary human myeloid cells, inflammasome activities in response to DNA transfection and cytosol-invasive microbes are mediated not by AIM2, but rather by NLRP3^68^. This human-specific DNA-induced NLRP3 activity occurs downstream of cGAS-STING activation^68^. The significance of this NLRP3-dependent response to DNA has not been examined in the context of infection with bacteria that do not directly invade the cytosol. We reasoned that if DNA release into the cytosol explains the phenotypes of *oatA* and *tarMS* mutant strains, then the activation of inflammasomes by *S. aureus* in human macrophages could be dependent on NLRP3 rather than AIM2. Consistent with this prediction, treatment of SA_Δ_*_oatA_*-infected hMDMs with the NLRP3 inhibitor MCC950 blocked IL-1β release (Fig 5E). Immunoblot analysis of infection supernatants revealed that cleaved IL-1β and cleaved caspase-1 were generated upon infection, and that production of these cleaved, activated proteins was prevented by treatment with MCC950 (Fig 5F). Treatment with the cGAS inhibitor G140 produced similar results, albeit to a lesser degree (Fig 5E,F). In contrast, MCC950, but not G140, blocked IL-1β release in response to the NLRP3 agonist nigericin, indicating that G140 does not block cGAS-independent NLRP3 activation.

These collective datasets in mouse- and human-derived cells support the idea that DNA is released into the cell during infection in a manner regulated by OatA and TarMS. To test this model directly, we quantified the amount of extrabacterial DNA (eDNA) in infected cells. To quantify eDNA within macrophages, we infected AIM2^-/-^ iBMDMs with *S. aureus* strains and lysed infected cells with RIPA buffer. Importantly, this lysis procedure did not lyse or trigger DNA release from *S. aureus* (Fig S3F). We then removed intact bacteria from infected cell lysates by centrifugation, isolated eDNA from these cleared iBMDM lysates, and performed qPCR to evaluate eDNA abundance. Using this assay, we found that Δ*oatA* strains released the most eDNA during infection, whereas Δ*tarMS* strains released the least (Fig 5G). These trends are consistent with our analyses of AIM2 and cGAS activation. OatA and TarMS therefore have competing and functionally important roles in controlling the release of bacterial eDNA (model schematic, Fig S4).

## DISCUSSION

Based on the data presented, we propose that activation of the cytosolic DNA sensors AIM2 and cGAS requires *S. aureus* WTA and is enhanced by WTA glycosylation. Several lines of evidence support this proposed model. First, deletion of the gene encoding the WTA glycosyltransferase TarM (or depletion of WTA by tunicamycin) disrupted AIM2 inflammasome activation in infected macrophages. Second, deletion of the gene encoding OatA increased both PGN-associated WTA abundance and AIM2 inflammasome activation. The opposing effects of WTA and OatA were observed at the functional level (by assessing IL-1β release from infected cells), biochemical level (by assessing caspase-1 and IL-1β cleavage), and single cell level (by assessing AIM2 inflammasome assembly in real time in infected cells). Third, the WTA glycosyltransferase activity of TarM was necessary for the function of TarM as a regulator of AIM2 activation. Fourth, the antagonistic phenotypes of WTA and OatA were not only observed for AIM2 activation but were also evident when examining bacterial DNA release into infected cells and cGAS activation. Fifth, WTA enhanced the activation of DNA-triggered innate immune receptors in human and mouse macrophages and during infections with clinical bacterial isolates. Based on these data, we propose that DNA-triggered inflammasomes, whether via AIM2 in mice or NLRP3 in humans, are common participants in the inflammatory response to nontoxigenic *S. aureus* infections.

While TarM and WTA have not previously been described as inflammasome or cGAS-STING regulators, prior research has explored the function of OatA in inflammasome responses to *S. aureus*. We can fully recapitulate prior findings that OatA prevents inflammasome activation during *S. aureus* infections of macrophages^22,33^. However, while OatA is known for its role in preventing PGN degradation by lysozyme *in vitro*^31,32,69^, our data indicate that lysozyme deficiency does not impact the inflammasome stimulatory activities of OatA-deficient *S. aureus*. Rather, we find that OatA restricts the abundance of WTA-modified PGN, thereby preventing AIM2 and cGAS activation.

One could interpret the finding that WTA (in particular, glycoWTA) promotes AIM2 and cGAS activation as evidence that this cell wall component is a newly defined PAMP. However, the compendium of evidence presented herein is inconsistent with this idea. Rather than operating as a PAMP, which is an intrinsically immunostimulatory entity (*e.g.* DNA), WTA controls the ability of DNA to operate as a PAMP. This hierarchy of bacterial factors, whereby one controls the actions of another during infection, is reminiscent of the emerging concept of bacterial effectors and meta-effectors^70–72^. Whereas effectors are virulence factors that directly manipulate the host to promote infection, meta-effectors are bacterial proteins that control the activities of the effectors within infected cells^70–72^. Drawing on this concept, we propose that WTA can be considered a meta-PAMP, which controls the intracellular activities of DNA, the actual PAMP. While PAMPs can be studied in isolation to identify their activities within infected cells, the immunomodulatory activities of meta-PAMPs cannot be revealed through their study in isolation. Rather, meta-PAMPs can only be identified through the study of the microorganism. As most PAMPs are buried in the bacterial cell (*e.g.* DNA, ribosomal RNA, peptidoglycan fragments and the lipid A component of LPS)^73^, it is possible that additional factors exist, akin to WTA, which facilitate PAMP release during infection. This concept and supporting evidence provide a mandate for additional microbe-centric (rather than PAMP-centric) analyses of the host-bacterial interaction.

## Methods

### Materials

*Escherichia coli* (*E. coli*) was grown in Miller LB broth (Sigma-Aldrich), *S. aureus* was grown in tryptic soy broth (Becton Dickinson/BD), and solid growth media was prepared using Difco agar (BD). BMDMs and iBMDMs were grown in high glucose DMEM (high glucose, glutamate (+), pyruvate (+); Thermo Fisher Scientific) supplemented with 10% FBS (R&D Systems) and 1x penicillin/streptomycin (Thermo Fisher Scientific) to generate complete DMEM (cDMEM). Human hMDMs were grown in RPMI (Thermo Fisher Scientific) supplemented with 10% FBS and 1x penicillin/streptomycin (cRPMI). Lysostaphin was obtained from Sigma-Aldrich. LPS O55:B5, nigericin, poly(dA:dT), MCC950, and zVAD-FMK were obtained from Invivogen. Human M-CSF was obtained from R&D. Uncoated mouse TNFα ELISA kits and LDH assay kits were obtained from Thermo Scientific. Lumit kits for mouse IL-1β and Wizard genomic DNA extraction kits were obtained from Promega. Uncoated mouse IP-10 ELISA kits were obtained from R&D. Antibodies used for immunoblotting were against mouse IL-1β (GeneTex), Gasdermin D (Cell Signaling), Caspase-1 (Adipogen), NLRP3 (Adipogen), AIM2 (cell Signaling), β-actin (BioLegend), and lysozyme (Abcam). Anti-human IL-1β antibodies were from GeneTex and Caspase-1 antibodies were from Adipogen. HRP-conjugated secondary antibodies were from Jackson ImmunoResearch. Cas9 and sgRNAs were from IDT.

### Clinical *S. aureus* isolates

Patient-derived *S. aureus* strains were obtained from the Brigham & Women’s Hospital clinical microbiology laboratory with approval by and following the guidelines of the Mass General Brigham Institutional Review Board (Protocol #2021P002127). Patients being actively treated for relapsed or recurrent *S. aureus* bacteremia at Brigham & Women’s Hospital were identified. Bacterial strains isolated from the blood of patients of interest were restreaked from blood agar plates generated by the clinical laboratory onto tryptic soy agar. Single colonies were picked into tryptic soy broth, grown overnight, and frozen as glycerol stocks until use.

### Data analysis and statistics

All experiments were performed at least 3 times to ensure reproducibility. For quantitative experiments, except where indicated otherwise data were pooled from 3 independent experiments, with 2-3 technical replicates performed for each condition during each experiment. Data were evaluated for statistical significance by Student’s *t*-test; ****: *p*<0.0001; ***: *p*<0.001; **: *p*<0.01; *: *p*<0.05; ns: not statistically significant. For qualitative experiments (*i.e.* immunoblots), a single experiment representative of at least 3 independent experiments is presented. Data was plotted and analyzed using Prism GraphPad software.

### Cloning, plasmids, and primers

Cloning was performed using *E. coli* strains DH5α and Stbl3. Plasmids pJC1111^74^ and pOS1-P*_lgt_*^50^ have been previously described and were generously provided by Drs. Victor J. Torres and Jean C. Lee, respectively. Plasmid pMSCV-IRES-Puro is a derivative of plasmid pMSCV-IRES-GFP and has been previously described by our group. Primers and gBlocks were obtained from IDT. PCR amplifications were performed using Phusion DNA polymerase (Thermo Fisher Scientific). Restriction digestion and ligation of PCR products to generate plasmids was performed using enzymes from New England Biolabs. GeneJet Plasmid Miniprep kits were obtained from Thermo Fisher, Nucleobond Xtra Midi kits were from Macherey-Nagel, and PCR Purification and Gel Extraction kits were from Qiagen. Sanger sequencing was performed by the Harvard Biopolymers Core Facility or by Azentaå. Whole-plasmid and bacterial whole genome sequencing were performed by Plasmidsaurus.

### Tissue culture and macrophage differentiation

To prepare primary murine BMDMs, marrow was harvested from the leg bones of wild-type C57BL6/J (The Jackson Laboratory, #000664), Casp1/11^-/-^ (#016621), NLRP3^-/-^ (#021302), or AIM2^-/-^ (#013144) mice. Briefly, mice were euthanized by CO_2_ asphyxiation and leg bones were dissected. Bones were cut at both ends and marrow was collected by centrifugation into a fresh microfuge tube. Red blood cells were lysed with ACK buffer (Gibco), and marrow was differentiated into primary BMDMs for 7 days in complete DMEM (cDMEM: 10% FBS, 1x penicillin/streptomycin) supplemented with 30% L929 supernatant. iBMDMs were grown in cDMEM and passaged every two to three days.

hMDMs were differentiated from peripheral blood mononuclear cells (PBMCs). Human K2-EDTA Buffy coats were obtained from BioIVT and mixed 1:1 with PBS supplemented with 2.5mM EDTA. PBMCs were isolated via density centrifugation by layering blood over Ficoll Paque Plus (Cytiva) and centrifuging at 800xg for 35 minutes without brake at room temperature. The PBMC layer was isolated, washed extensively in MACS buffer (PBS supplemented with 2.5mM EDTA, 2% FBS, and 1x penicillin/streptomycin), and red blood cells were lysed with ACK buffer. Monocytes were isolated using magnetic anti-CD14 beads (Miltenyi) and then differentiated into hMDMs by plating on tissue culture-treated plates for 6 days in cRPMI supplemented with 30 ng/mL human M-CSF (R&D). On days 2 and 4 of differentiation, half of the tissue culture media was removed; nonadherent cells were pelleted by centrifugation, resuspended in an equal volume of fresh cRPMI supplemented with 30 ng/mL M-CSF, and added back to culture plates. On day 6, differentiated hMDMs were lifted from plates using macrophage detachment solution as per the manufacturer’s instructions (PromoCell) and plated in cRPMI without M-CSF. Infections were performed on day 7.

### Bacterial culture

Bacterial strains of interest were streaked onto either tryptic soy agar (*S. aureus*) or LB agar (*E. coli*) plates from frozen glycerol stocks. Individual colonies were picked into TSB or LB supplemented with antibiotics as needed and grown overnight at a 45° angle with shaking at 37°C. For *E. coli*, selection was achieved with ampicillin (100 µg/mL), chloramphenicol (25 µg/mL), or kanamycin (50 µg/mL). For *S. aureus*, selection was achieved with chloramphenicol (10 µg/mL), kanamycin (25 µg/mL), cadmium (0.1 mM), or erythromycin (5 µg/mL). To evaluate hemolytic activity, indicated strains were streaked from frozen glycerol stocks onto TSA-5% sheep’s blood agar (BD) and grown overnight at 37°C.

### Preparation of electrocompetent S. aureus

A single colony of the strain to be transformed was picked into 3 mL TSB and grown overnight. The following morning, the overnight culture was diluted 1:100 into 30 mL fresh TSB and grown at 37°C for 3 hours. Bacteria were then chilled on ice, pelleted at 4122 xg for 8 minutes, washed twice with 30 mL and then once with 15 mL ice-cold sterile 10% glycerol. Bacteria were resuspended in 3 mL of cold 10% glycerol and flash frozen in 50 µL aliquots. Frozen electrocompetent bacteria were thawed on ice, mixed with the indicated plasmid DNA, and then electroporated using cuvettes with a 0.2 cm electrode gap (Bio-Rad) with the following conditions: 1800 V, 10 µF, 600 Ω. 1 mL of fresh TSB was immediately added to electroporated bacteria, which were incubated with shaking at 37°C for 1.5 hours and then plated on selective media to identify transformants.

### Construction of S. aureus strains

*S. aureus* strain SA113 (SA_WT_) was generously provided by Dr. Jean C. Lee. A strain lacking OatA (SA_Δ_*_oatA_*) was generated by packaging a deletion/insertion cassette into Phage 80α using the previously described Δ*oatA*::*kan* mutant^22,31^ as a donor strain, transducing SA_WT_, and selecting for mutants on kanamycin. *oatA* and *tarM* complementation was achieved by cloning the indicated gene from SA_WT_ along with ∼200 base pairs of upstream promoter sequence into the plasmid pJC1111^74^. Empty pJC1111, pJC1111-*oatA*, or pJC1111-*tarM* was transformed into recombineering strain RN9011^74^ and integrants were selected using cadmium. DNA from integrants was then packaged into Phage 80α and used to transduce either SA_WT_, SA_Δ_*_oatA_*, or transposon mutant 3H4, followed by selection on cadmium.

Unmarked deletions of *tarM*, *tarS, graRS*, and *oatA* (SA_Δ_*_tarM_*, SA_Δ_*_tarS_*, SA_Δ_*_tarMS_*, SA_Δ_*_graRS_*, SA_Δ_*_oatA_*) were generated using the plasmid pIMAY*^75^. Briefly, 1kb of sequence each upstream and downstream of the target gene was cloned into pIMAY* to generate knockout (KO) plasmids; the first and last 3-5 codons of each gene were retained. Each KO plasmid was used to transform *E. coli* strain IM08B^76^, and IM08B-derived plasmid was then electroporated into electrocompetent *S. aureus*. Transformants were selected by growth on chloramphenicol at permissive temperature (29°C). Integration was achieved by passaging into fresh chloramphenicol-supplemented media and growing overnight at non-permissive temperature (37°C), followed by plating on chloramphenicol-supplemented TSA at 37°C. Large bacterial colonies representing integrants were inoculated into TBS lacking antibiotics and passaged twice per day at 29°C for a total of 6 passages, and then serially diluted in TSB and plated on LB agar supplemented with 20 mM para-chlorophenylalanine (Thermo Fisher Scientific) at 37°C. Large colonies were patched onto TSA either with or without chloramphenicol, and chloramphenicol-susceptible isolates representing putative deletion mutants were streaked out for single colonies. Genomic DNA was prepared from each potential mutant, and PCR using primers that anneal outside of the KO plasmid sequence was performed to identify successful deletions. gDNA from mutant strains was sent for whole genome sequencing for unbiased confirmation of the target deletions. Mutants were subsequently stored as glycerol stocks for use in further experiments.

Restoration of glycosyltransferase expression and expression of superfolder GFP was accomplished by cloning *tarS* (from SA113), *tarM* (from SA113), *tarP* (sequence from strain N315 was synthesized as a gBlock), or superfolder GFP from plasmid pSR1037^77^ into the multicloning site of plasmid pOS1 downstream of the constitutively active P*_lgt_* promoter^50^ *tarM*_E403A_ was generated from WT *tarM* by overlap extension PCR. Plasmids were used to transform and subsequently purified from *E. coli* strain IM08B, electroporated into SA_Δ_*_oatA_*_/Δ_*_tarMS_*, and selected by plating on TSA supplemented with chloramphenicol. Strains transformed with pOS1 were maintained in 10 µg/mL chloramphenicol for all subsequent growth.

### Bacterial infection and inflammasome activator treatment of macrophages

For infections analyzed by Lumit, ELISA, and LDH assay, the day prior to infection either 100,000 cells/well primary BMDMs, 100,000 cell/well primary hMDMs, or 50,000 cells/well iBMDMs were plated in triplicate in a 96-well plate to yield 100,000 cells per well the following day. On the day of infection, plating media was exchanged for priming media (DMEM/FBS or RPMI/FBS with or without 1 µg/mL LPS) and incubated at 37°C for 3 hours. Overnight cultures of bacteria were washed twice with cold PBS, resuspended in DMEM/FBS, and diluted to achieve an MOI of 10. For all infection experiments, *S. aureus* strains were serially diluted and plated in triplicate on TSA to enumerate CFU. Primed cells were placed on ice and priming media was removed and replaced with infection media. Plates were centrifuged at 500xg for 5 minutes at 4°C to coordinate phagocytosis and then placed at 37°C to initiate infection. 30 minutes post-infection, infection media was replaced with DMEM supplemented with 10% FBS and 20 µg/mL lysostaphin. 4 hours post-infection, plates were centrifuged at 400xg for 3 minutes and supernatants were collected for further analysis. Lumit, ELISA, and LDH assays were performed as per the manufacturers’ protocols. Where indicated, inhibitors were added to tissue culture media 30 minutes prior to infection and maintained for the remainder of the experiment. For experiments utilizing cytochalasin D, bafilomycin A1, and ammonium chloride, infection media was left in place and lysostaphin was not added to avoid removing or lysing non-phagocytosed bacteria.

For infections analyzed by SDS-PAGE and immunoblotting, the day prior to infection 500,000 BMDMs or hMDMs were plated per well in a 24-well plate. On the day of infection, cells were primed and infected as above. 30 minutes post-infection, infection media was removed, cells were washed once with PBS, and OptiMEM supplemented with 20µg/mL lysostaphin was added. 4 hours post-infection, plates were centrifuged at 400xg for 3 min and supernatants were removed. Cells were washed once with cold PBS and then lysed by addition of 200 µL reducing denaturing sample buffer followed by homogenization through a syringe with a 25G needle. An aliquot of supernatant was mixed with reducing denaturing sample buffer; for hMDMs, the remainder of supernatant was incubated rotating with a biotinylated anti-IL-1β antibody (R&D) and Neutravidin resin (Thermo Fisher) overnight at 4°C, washed three times with cold PBS, and released from resin by mixing with reducing denaturing sample buffer. Supernatants, immunoprecipitation eluates, and lysates were boiled for 5 minutes, separated by SDS-PAGE on a 12% polyacrylamide gel, and transferred to a PVDF membrane. Membranes were blocked in tris-buffered saline with 0.1% Tween-20 (TBST) containing 3% milk and incubated overnight at 4°C with the indicated primary antibody at a 1:1000 dilution prepared in TBST-1% bovine serum albumin (BSA). Membranes were washed with TBST, incubated with an HRP-conjugated secondary antibody prepared at a 1:5000 dilution in TBST-3% milk, washed with TBST, and developed using SuperSignal West Pico Plus or Femto chemiluminescence kits (Thermo Scientific). Images were obtained using a Bio-Rad ChemiDoc imaging system.

For experiments utilizing nigericin, BMDMs were primed with LPS for 3 hours as above, and nigericin was added to the media at a final concentration of 10 µM for 2 hours. For experiments utilizing poly(dA:dT), BMDMs were primed and then transfection was performed using Lipofectamine 2000. Briefly, 1 µg poly(dA:dT) per 100,000 cells was mixed with OptiMEM; for each 1 µg of poly(dA:dT), 2 µL Lipofectamine 2000 was mixed with OptiMEM. Lipofectamine and poly(dA:dT) mixtures were then combined, incubated for 30 minutes at room temperature, and then added to DMEM/FBS to reach 1 µg poly(dA:dT) per 200 µL media. Priming media was removed from BMDMs and replaced with poly(dA:dT) transfection mixture; cells were incubated for 2 hours prior to collection.

### Generation of iBMDM knockout lines and AIM2-mScarlet iBMDMs

iBMDM knockouts were generated using CRISPR-Cas9 ribonucleoprotein complexes (RNPs). RNPs were generated by mixing recombinant Cas9 with a sgRNA targeting either AIM2 or a combination of sgRNAs targeting Lyz1 and Lyz2. RNPs were electroporated into 1 x 10^6^ WT iBMDMs (Neon Transfection System, Thermo Fisher) and plated in a 6 well plate to recover overnight. The following day, cells were resuspended, diluted in cDMEM, and plated to separate single cell clones in 96 well plates. Genomic DNA and whole cell lysates were prepared from single cells clones and were analyzed by PCR using primers flanking the sgRNA target site to identify clones in which all copies of target genes were disrupted and by immunoblotting to confirm loss of the target gene product.

To generate AIM2-mScarlet-producing cells, a gBlock was synthesized encoding the mouse AIM2 gene fused to a C-terminal flexible linker followed by the mScarlet gene and was cloned into the retroviral vector pMSCV-IRES-Puro. Either empty or AIM2-mScarlet plasmid was used to transfect HEK293T cells along with packaging plasmid pCL-Eco and pVSV-G using Lipofectamine 2000 (Invitrogen). HEK293T supernatants containing retrovirus were passed through a 0.45 µM filter, supplemented 1:2000 with polybrene, and then used to transduce WT or AIM2^-/-^ iBMDMs with centrifugation at 1250xg for 1 hour at 30°C. 48 hours after transduction, tissue culture media was supplemented with 5 µg/mL puromycin and cells were grown and passaged for one week to select for transductants; puromycin was maintained for all further growth. To achieve a population with homogenous AIM2-mScarlet expression, transductants were diluted in cDMEM and plated in 96 well plates to achieve single cell clones. Individual clones were analyzed by immunoblotting for AIM2-mScarlet production and by SA_Δ_*_oatA_* infection to confirm rescue of AIM2 activity.

### SAΔ_oatA_ forward genetic screen

A transposon mutant library was generated in SA_Δ_*_oatA_* using a previously described system. SA_Δ_*_oatA_* was transformed sequentially with the plasmid pBursa encoding the mariner-based *bursa aurealis* transposon^40^ and plasmid pMG020^78^ (generously provided by Dr. Anthony R. Richardson) encoding Himar1 transposase under control of the constitutively active P*_lgt_* promoter. Double transformants were inoculated into TSB and then plated for single colonies representing individual transposon insertion clones.

For screening infections, WT iBMDMs were plated the day prior to infection at a density of 12,500 cells/well in white-walled tissue culture-treated 384-well plates. On the day of infection, iBMDMs were primed for 3 hours with 1 µg/mL LPS. Individual bacterial colonies were inoculated into DMEM/FBS and vortexed; an aliquot of this inoculum was used to directly infect primed iBMDMs and the remainder was grown overnight in a 96 well plate at 37°C. 30 minutes post-infection, infection media was supplemented with lysostaphin to a final concentration of 20 µg/mL. 4 hours post-infection, an *in situ* Lumit IL-1β assay was performed. Wells in which IL-1β release was reduced by 50% or more were considered putative hits. Bacterial isolates corresponding to these wells were streaked onto TSA to obtain single colonies, and single colony isolates were then grown overnight in TSB; liquid overnight cultures were used to infect iBMDMs for phenotype validation. Validated single colony isolates that triggered reduced IL-1β release during infection of iBMDMs were considered hits and were stored as glycerol stocks for further experiments. The identification of transposon insertion sites and confirmation of a single insertion per strain was accomplished by purification of genomic DNA using the Promega Wizard kit and whole genome sequencing by Plasmidsaurus.

### WTA purification, PAGE, and silver staining

WTA was purified from indicated *S. aureus* strains using published methods^79^. Briefly, *S. aureus* was grown overnight in TSB with or without addition of 0.4 µg/mL tunicamycin. Overnight cultures were washed sequentially with TSB and Buffer 1 (50 mM MES pH 6.5), followed by resuspension in Buffer 2 (50 mM MES pH 6.5, 4% SDS). Bacteria were submerged in a boiling water bath for one hour, cooled to room temperature, and cell walls were sequentially pelleted and washed with Buffer 1, Buffer 2, Buffer 3 (50 mM MES pH 6.5, 2% sodium chloride), and Buffer 1. Washed pellets were resuspended in Digestion Buffer (20 mM Tris pH 8.0, 0.5% SDS) supplemented with 20 µg/mL Proteinase K and incubated with shaking at 50°C for 4 hours to generate sacculus. Sacculus was pelleted and washed with Buffer 3 and then three times with water. To hydrolyze WTA, sacculus was resuspended in 0.1M sodium hydroxide and incubated with shaking overnight at room temperature. Sacculus was pelleted and supernatant containing WTA was removed; sodium hydroxide was neutralized by addition of 1M Tris-HCl pH 7.8.

WTA extracts were mixed 3:1 with 4x sample buffer (50 mM Tricine, 50 mM Tris pH 8.2, 50% glycerol, trace bromophenol blue) and separated on a Novex 16% Tricine polyacrylamide gel (Invitrogen) at 125V for 90 minutes in non-denaturing running buffer (100 mM Tris, 100 mM Tricine, pH 8.0) at 4°C. Gels were silver stained using the ProteoSilver Plus kit (Sigma-Aldrich) and imaged using a ChemiDoc imaging system.

### Quantitative real-time PCR to measure expression of S. aureus genes

Indicated *S. aureus* strains were grown overnight in TSB, diluted 1:100 in fresh TSB, and grown to either mid-log (OD_600_ 1-2), late log/early stationary (OD_600_ 5-6), or deep stationary (overnight culture) phase. Upon reaching target OD_600_, cultures were cooled on ice, pelleted, resuspended in lysostaphin buffer (10 mM Tris pH 8.2, 0.1 mM EDTA, 50 µg/mL lysostaphin), and incubated at 37°C for 30 mins. RNA was purified using PureLink RNA mini kit (Invitrogen), and DNA was removed by treatment with DNase I (NEB). cDNA was generated from total RNA using the MultiScribe High-Capacity cDNA Reverse Transcription Kit (ThermoFisher Scientific). Reaction mixtures were diluted 1:10 with water and used as a template for qPCR reactions with iTaq Universal SYBR Green Supermix (Bio-Rad) with primers targeting the coding sequence of *S. aureus* 16S RNA, *tarO*, *tarM*, or *tarS*.

### Measurement of internalization of S. aureus by macrophages

To measure internalization of *S. aureus*, WT BMDMs were plated at a density of 250,000 cells/well onto a glass coverslip in a 24 well plate. The following day, the indicated bacterial strains were prepared as above and BMDMs were infected at an MOI of 1. One hour post-infection, cells were fixed by adding paraformaldehyde to the culture media at a final concentration of 4% and incubated at room temperature for 15 minutes. Fixed cells were washed 4 times in cold PBS and incubated in PBS overnight at 4°C. Coverslips were blocked with PBS supplemented with 0.2% BSA, and then extracellular *S. aureus* was stained green by incubating with a rabbit anti-*S. aureus* antibody prepared at a 1:100 dilution in blocking buffer, washing with PBS, and then incubating with an AlexaFluor488-labeled anti-rabbit IgG antibody prepared at a dilution of 1:250 in blocking buffer. Coverslips were washed with PBS and then fixed again with 4% PFA to stabilize staining of extracellular bacteria. Coverslips were washed with PBS and residual PFA was quenched with PBS supplemented with 50 mM ammonium chloride. Coverslips were then incubated with permeabilization buffer (PBS supplemented with 0.2% BSA and 0.05% saponin). Total *S. aureus* was stained red by incubating with the same rabbit anti-S*. aureus* antibody prepared at a 1:100 dilution in permeabilization buffer followed by incubation with an AlexaFluor594-conjugated anti-rabbit IgG antibody. Coverslips were washed with PBS, rinsed once in water, and mounted onto glass slides using ProLong Glass Antifade mountant.

Imaging of fixed samples was performed on a Zeiss LSM 880 point-scanning confocal microscope with fast Airyscan controlled with Zeiss Zen Blue v.2.3SP1 software. Imaging was performed using a Zeiss Plan Apochromat x40/1.3 oil-immersion objective. 3 fields of view were imaged for each sample; 9 Z-stacks were obtained for each position covering a total range of 6.7 µm. Microscope settings were as follows: bidirectional scanning, 2x frame averaging, 16-bit depth, ROI 5600 x 5600 px with a pixel size of 67 nm. Following imaging, the acquired image stacks were mean intensity projected, normalized and median filtered to remove salt-and-pepper noise. All bacteria were segmented using the “BactFluorOmni” neural network as implemented in Omnipose c1.0.6^80^. The resulting bacteria labels were filtered for labels >0.25 µm^2^ to remove artifacts. Next, the bacteria labels were used to determine the average intensity of each label (AlexaFluor488 for extracellular bacteria; AlexaFluor594 for total bacteria) and the labels categorized into extracellular and total based on the calculated intensity (considered extracellular if avg. fluorescence intensity >10X background of unstained bacteria). Subsequently, the proportion of intracellular and extracellular bacteria was calculated. Next, cells were segmented using the “cpsam” neural network as implemented in Cellpose v4.0.4^81^ and cell and intracellular (stain-negative) bacteria labels used to calculate the %infected cells and infection levels (number of bacteria per infected cell) using labeling arithmetic.

### Live cell imaging of AIM2-mScarlet iBMDMs

For quantification of AIM2 specks and SYTOX Green uptake, on the day prior to infection, 250,000 AIM2^-/-^-AIM2-mScarlet iBMDMs were plated in cDMEM supplemented with 5 µg/mL puromycin in a 24-well glass bottom plate (MatTek #P24G-1.5-10-F) and incubated overnight at 37°C. On the day of infection, bacteria were prepared as above but resuspended in imaging media (Phenol red-free DMEM supplemented with glutamate, pyruvate, 10% FBS, and 20 mM HEPES) supplemented with 0.5 µM SYTOX Green (Invitrogen) at a target MOI of 10. Plating media was replaced with 400 µL cold infection mixtures and cells were centrifuged at 500xg for 5 minutes to coordinate phagocytosis, and cells were kept on ice and transported to a microscope with stage pre-warmed to 37°C. Infection time = 0 was considered the moment that the cells were placed on the pre-warmed stage. 30 minutes post-infection, 100 µL of prewarmed imaging media supplemented with lysostaphin was added to give a final concentration of 20 µg/mL. For direct stimulation of AIM2, Lipofectamine 2000 was used to transfect 5 µg of poly(dA:dT). For direct stimulation of NLRP3, cells were primed with 1 µg/mL LPS for 3 hours followed by the addition of nigericin to a final concentration of 10 µM.

Imaging of live samples was performed on a Nikon Ti2-E inverted widefield microscope using a Plan Apo λ 20x/0.75 DIC N2 air objective. 3 fields of view were obtained for each sample. Microscope settings were as follows: 16-bit depth, ROI 1952 x 1952 px with a pixel size of 216 nm. Images were obtained every minute for 4 hours. Recorded image stacks were corrected for photobleaching using histogram matching^82^ as implemented in ImageJ/Fiji. Cells were segmented using the “cpsam” neural network as implemented in Cellpose v4.0.4^81^, and the resulting cell labels counted following exclusion of labels >2 SD smaller than the average label (removing debris/artefact labels resulting from noise) and of labels touching the edge of the field of view (incompletely captured cells). Quantification of AIM2 specks and SYTOX Green positive cells at each position was performed manually using the ImageJ Cell Counter plugin and converted to a percentage by dividing the number of specks or SYTOX Green positive cells by the above calculated total cell number at that position.

For measurements of distance between AIM2 specks and bacteria, on the day prior to infection, 1.5 million AIM2^-/-^+AIM2-mScarlet iBMDMs were plated in cDMEM supplemented with 5 µg/mL puromycin on a glass bottom 35 mm dish (MatTek #P35G-1.5-14-C) and incubated overnight at 37°C. 15 minutes prior to infection, media was supplemented with zVAD-FMK. SA_Δ_*_oatA_* was prepared as above and resuspended in 1.5 mL cold imaging media for a target MOI of 10. Plating media was removed from iBMDMs and replaced with cold infection media. Cells were centrifuged at 500xg for 5 minutes at 4°C to coordinate phagocytosis, and cells were kept on ice and transported to a microscope with stage pre-warmed to 37°C. Infection time = 0 was considered the moment that the cells were placed on the pre-warmed stage. 30 minutes post-infection, 0.5 mL of prewarmed imaging media supplemented with lysostaphin was added to give a final concentration of 20 µg/mL.

Imaging was performed on a Nikon Ti2-E inverted widefield microscope using a Plan Apo λ 20x/0.75 DIC N2 air objective. 6 fields of view were obtained for each sample; 5 Z-stacks were obtained for each position covering a total range of 10 µm. Microscope settings were as follows: 16-bit depth, ROI 1952 x 1952 px with a pixel size of 216 nm. Images were obtained every minute for 3 hours. Recorded image stacks were corrected for photobleaching using histogram matching^82^ as implemented in ImageJ/Fiji. Identification of AIM2 specks and measurement of distance to the nearest green bacterium was performed manually using Fiji/ImageJ.

### Measurement of bacterial survival within macrophages

Survival of intracellular bacteria was evaluated using a published protocol with slight modifications^83^. WT BMDMs were plated in 96 well plates at a density of 100,000 cells/well and infected as above with the indicated *S. aureus* strains at an MOI of 1; lysostaphin was not added. At the indicated time points post-infection, cells were washed three times with PBS and lysed in 50 µL PBS-1% saponin for 15 minutes at 37°C. Bacteria released from lysed BMDMs were serially diluted and plated on TSA to enumerate surviving CFU.

### Quantification of bacterial DNA inside infected macrophages

On the day prior to infection, AIM2^-/-^ iBMDMs were plated in 12 well plates at a density of 500,000 cells/well to give 1,000,000 cells/well at the time of infection. On the day of infection, iBMDMs were infected as described above with the indicated *S. aureus* strains, and after 30 minutes infection media was exchanged for fresh media supplemented with 20 µg/mL lysostaphin. 3 hours post-infection cells were washed three times with cold PBS and lysed by addition of 500 µL RIPA buffer (50 mM Tris pH 7.4, 150 mM sodium chloride, 1% NP-40, 0.5% sodium deoxycholate, 0.1% SDS) and rotating for 15 minutes at 4°C. Intact bacteria were pelleted by centrifugation, and supernatants were subjected to phenol-chloroform extraction to isolate solubilized DNA. RNA was removed by adding 1 µL RNase A and incubating at 65°C for 1 hour, followed by another round of phenol-chloroform extraction. Extracted DNA was precipitated by addition of 0.1 volume of 3M sodium acetate pH 5.2, 2 µL pellet paint coprecipitant (Millipore), and 2 volumes 100% ethanol, and pelleted by centrifugation. Pelleted DNA was washed once with 70% ethanol and DNA pellets were air dried before being resuspended in molecular grade water. qPCR was performed with iTaq Universal SYBR Green Supermix using primers targeting the coding sequence for *S. aureus* 16S RNA and mouse β-actin. 16S levels were normalized to β-actin. To quantify bacterial DNA, purified *S. aureus* gDNA was serially diluted to generate a standard curve.

### Infection of cGAS reporter iBMDMs

On the day prior to infection, MyD88/TRIF/STING^-/-^ iBMDMs transduced with either empty vector or pLenti-STING_R138H_ were plated in 96 well plates at a density of 50,000 cells/well for IP-10 ELISAs (to give 100,000 cells/well at time of infection) or in 24 well plates at a density of 250,000 cells/well for immunoblotting (to give 500,000 cells/well at time of infection). The following day, cells were infected with the indicated *S. aureus* strains, and 30 minutes post-infection media was replaced by fresh media supplemented with lysostaphin. 4 hours post-infection, cells were centrifuged, supernatants were removed, and cells were washed with cold PBS and lysed with reducing denaturing sample buffer. Supernatants were analyzed by IP-10 ELISA, and lysates were separated on a 12% polyacrylamide gel, transferred to a PVDF membrane, and immunoblotted for the indicated proteins.

## Acknowledgements

We thank members of the Kagan lab for helpful discussions that shaped this work. We thank Drs. Victor J. Torres, Jean C. Lee, and Anthony R. Richardson for providing bacterial strains and plasmids essential to this work. We appreciate important discussions and suggestions from Drs. Marcia Goldberg, Suzanne Walker, Charles L. Evavold, and Josue Flores-Kim. We thank Dr. Michael Anderson of the Boston Children’s Hospital Harvard Digestive Diseases Center core facility for assistance with the confocal microscope. We thank Dr. Paula Montero-Llopis and the staff of the Harvard Medical School Microscopy Resources on the North Quad (MicRoN) facility for providing equipment and assistance for widefield fluorescence microscopy studies. This work was funded by the Damon-Runyon Cancer Research Foundation grant DRG-123-24 (to JBJ), NIAID grants T32 AI7061 and K08 AI193234 (to JBJ), R01 AI167993 and R37 AI116550 (to JCK). DF was supported by a Human Frontier Science Program (HFSP) long-term postdoctoral fellowship (LT0006/2022-L) and an EMBO postdoctoral fellowship (ALTF 491-2022). Some images were prepared using BioRender.

## Author contributions

JBJ and JCK conceived of the study. JBJ and AT performed experiments. SAR generated cell lines. DF analyzed and contributed to the design of microscopy studies. JBJ and JCK prepared the manuscript.

---

**Figure S1.**
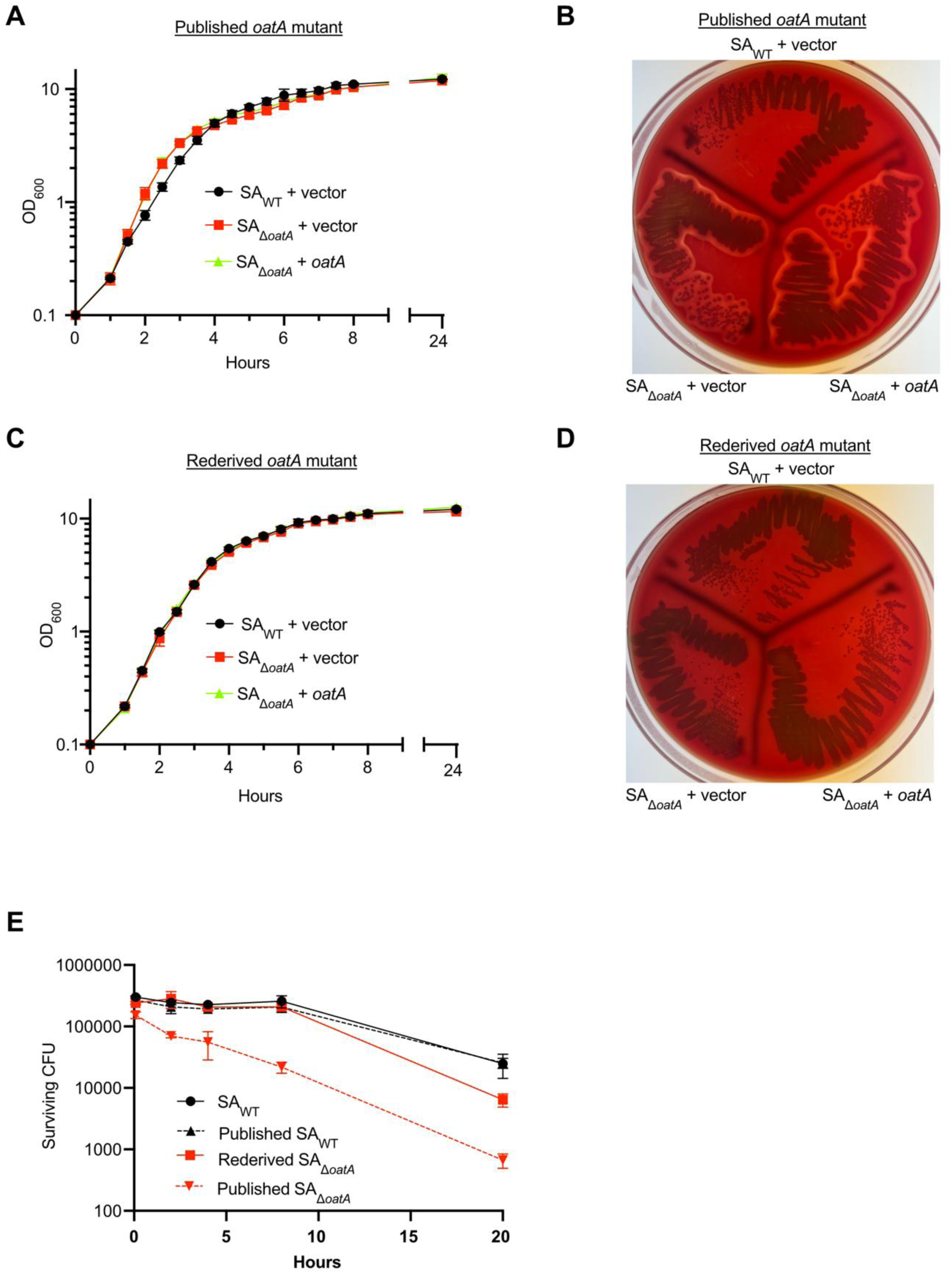
(A,C) The indicated *S. aureus* strains were diluted to OD_600_ = 0.1 in tryptic soy broth (TSB) and grown at 37°C. OD_600_ was checked after 1 hour and then every 30 minutes. (B,D) The indicated *S. aureus* strains were streaked onto TSA-5% sheep’s blood agar and grown overnight at 37°C. (E) Unprimed WT BMDMs were infected with the indicated *S. aureus* strains at an MOI of 1, and lysostaphin was added 30 minutes post-infection. At the indicated timepoints, cells were washed with PBS, lysed with PBS-1% saponin to release intracellular bacteria, serially diluted, and plated on solid media to enumerate surviving intracellular CFU.

**Figure S2.**
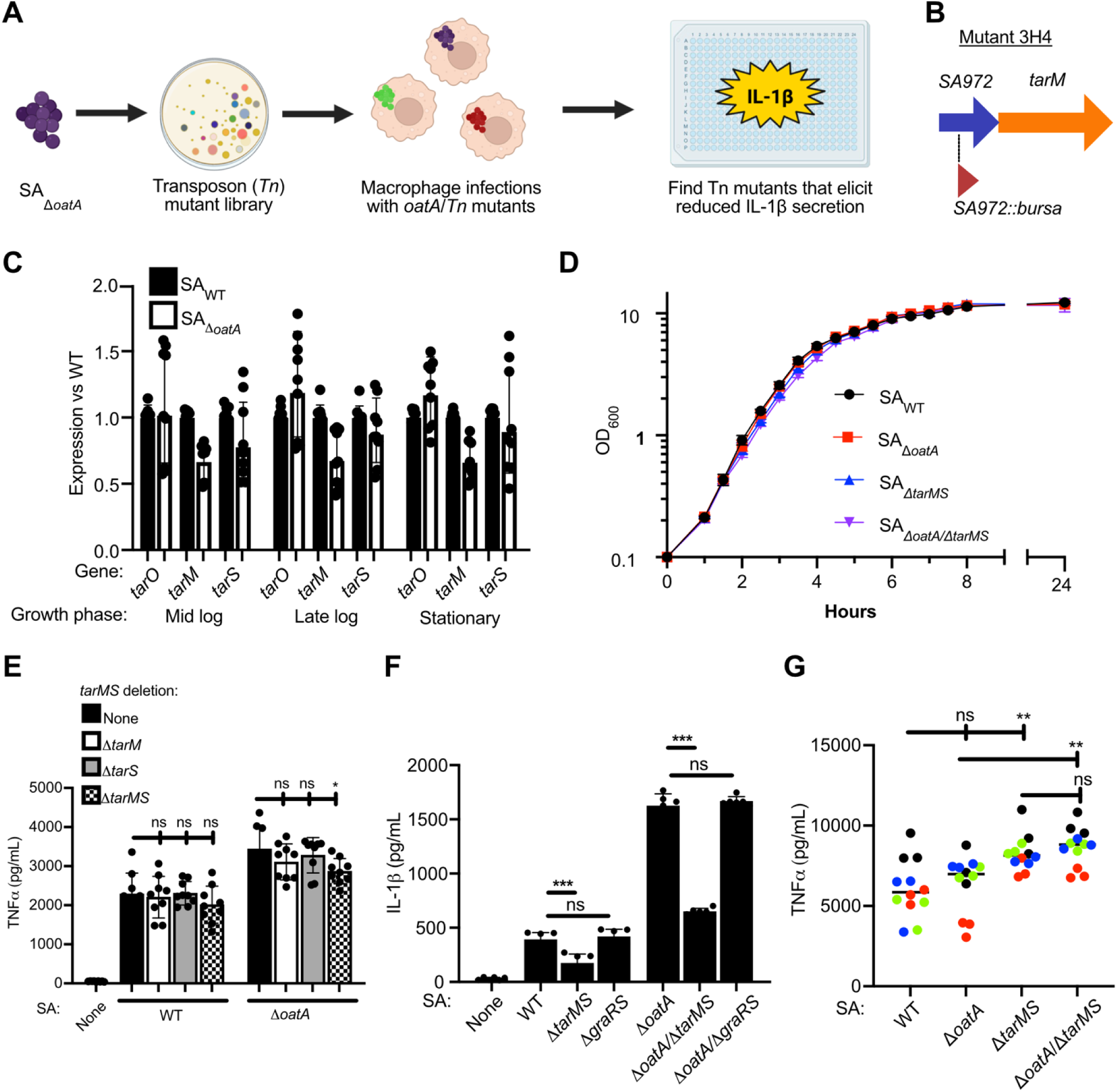
(A) Cartoon depicting bacterial suppressor screen design. SA_Δ_*_oatA_* was subjected to genome-wide mutagenesis by expressing Himar1 transposase in the presence of a plasmid encoding the *bursa aurealis* transposon. Individual bacterial colonies representing transposon insertion mutants were picked into tissue culture media, which was used to directly inoculate LPS-primed WT iBMDMs. After 4 hours, supernatants were analyzed in situ by Lumit assay for IL-1β. Mutants that triggered less IL-1β release than the parental SA_Δ_*_oatA_* strain were validated by repeat infections and had whole genomes sequenced to identify the site of transposon insertion. (B) Site of transposon insertion in mutant 3H4. The *bursa aurealis* transposon was located within the SA972 gene, which is encoded upstream of *tarM* within a two-gene operon. (C) The indicated *S. aureus* strains were diluted to OD_600_ = 0.1 in TSB and grown at 37°C until reaching mid-log (OD_600_ ∼1-2), late-log (OD_600_ ∼4-5), or stationary phase (overnight). Bacteria were pelleted, RNA was purified and used to generate cDNA. qPCR was performed using primers specific to *S. aureus* 16S, *tarO*, *tarM*, or *tarS*. Expression of each gene was normalized to 16S and plotted as a ratio of SA_Δ_*_oatA_* to SA_WT_. (D) The indicated *S. aureus* strains were diluted to OD_600_ = 0.1 in TSB and grown at 37°C. OD_600_ was checked after 1 hour and then every 30 minutes. (E,F) Unprimed (E) or LPS-primed (F) WT BMDMs were infected with the indicated *S. aureus* strains at a MOI of 10. 4 hpi, supernatants were collected and analyzed by Lumit assay for IL-1β. (G) Unprimed hMDMs were infected with the indicated *S. aureus* strains at an MOI of 10. 4 hpi, supernatants were collected and analyzed by ELISA for TNFα.

**Figure S3.**
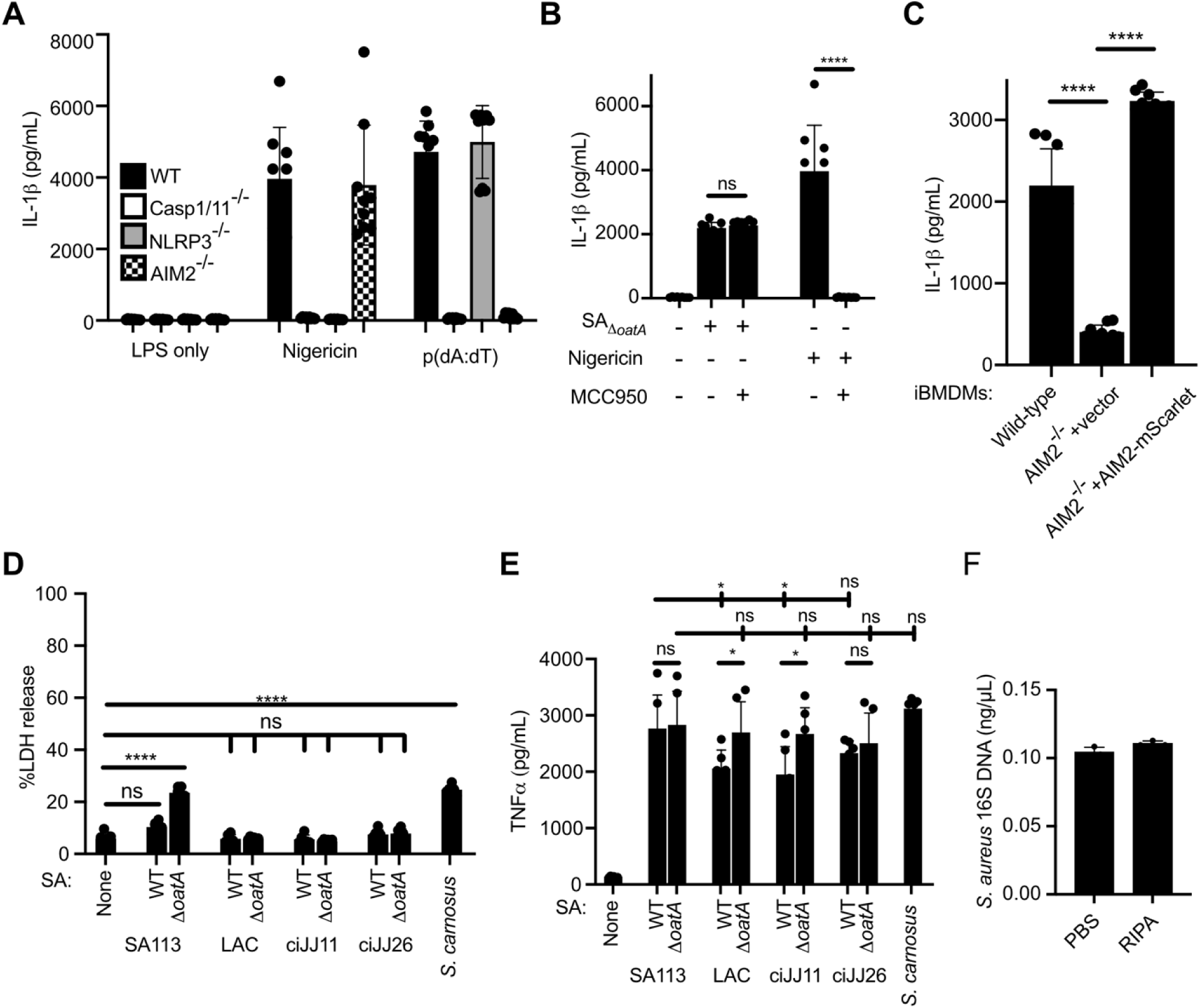
(A) WT BMDMs were primed with LPS and either infected with SA_Δ_*_oatA_* at an MOI of 10 for 4 hours or treated with 10 µM nigericin for 2 hours. Where indicated, cells were pretreated for 30 min and subsequently maintained in 5 µM MCC950. Supernatants were collected and analyzed by Lumit assay for IL-1β. (B) The indicated iBMDM lines were infected with SA_Δ_*_oatA_* at an MOI of 10 as above. 4 hpi, supernatants were collected and analyzed by Lumit assay for IL-1β. (C) Supernatants from Figure 4H were analyzed by LDH assay. (D) Unprimed WT BMDMs were infected with the indicated staphylococcal strains at an MOI of 10. 4 hpi, supernatants were collected and analyzed by ELISA for TNFα. (E) SA_Δ_*_oatA_* was incubated for 10 minutes at 4°C in either PBS or RIPA buffer, pelleted by centrifugation, and DNA was purified from supernatants. Bacterial DNA was quantified by qPCR with primers specific for the gene encoding 16S RNA. Data are representative of 3 independent experiments.

**Figure S4.**
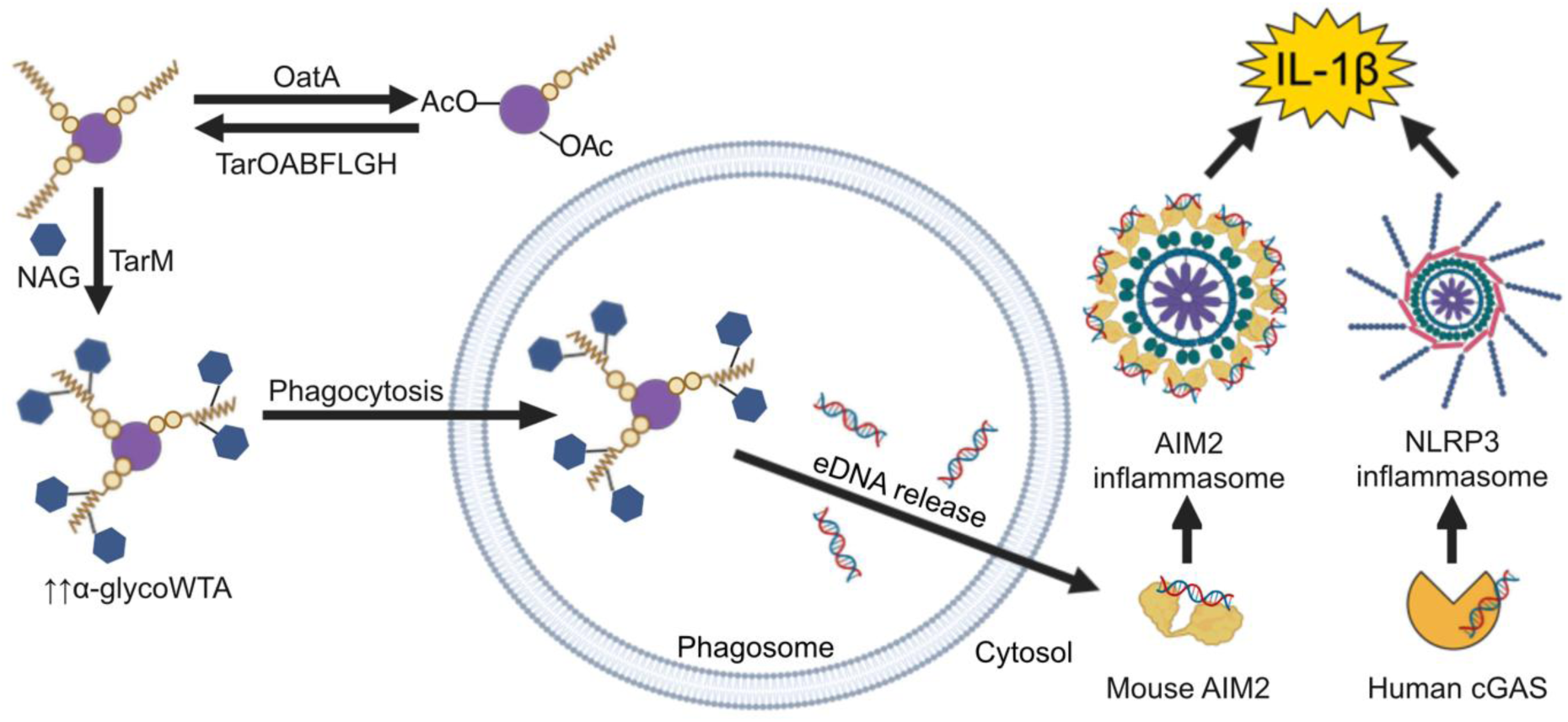
Working model of the regulation of inflammasomes by competing *S. aureus* cell wall modifications.

**Supplemental table 1.** Bacterial strains used in this study.

**Supplemental table 2.** Plasmids used in this study.

**Supplemental table 3.** Primers used in this study.

**Supplemental table 4.** CRISPR-Cas9 sgRNAs used in this study.

