## Supplemental Table 1 for "Wall teichoic acid is required for DNA-triggered innate immune receptor activation by *Staphylococcus aureus*"

**Supplemental Table 1. Bacterial strains used in this work**

| Species | Strain name | Background strain | Genotype | Plasmid | Source | Notes |
|---|---|---|---|---|---|---|
| *Escherichia coli* | | DH5a | *fhuA2Δ(argF-lacZ)U169 phoA glnV44 Φ80Δ(lacZ)M15 gyrA96 recA1 relA1 endA1 thi-1 hsdR17* | | New England Biolabs | Used for most cloning |
| | | Stbl3 | $F^-$ *mcr*B *mrr hsd*S20($r_B$–, $m_B$–) *rec*A13 *sup*E44 *ara*-14 *gal*K2 *lac*Y1 *pro*A2 *rps*L20(StrR) *xyl* -5 λ–*leu mtl*-1 | | Thermo Fisher Scientific | Used for cloning into pMSCV-IRES-puro |
| | | IM08B | SA08BΩ$P_{N25}$-*hsdS* (CC8-1) (SAUSA300_0406) of NRS384 integrated between the *essQ* and *cspB* genes | | Monk et al, 2015; Provided by Victor J. Torres | Used for passaging plasmids prior to electroporation into *S. aureus* |
| *Staphylococcus aureus* | SaJJ19 | SA113 | WT | | Jean C. Lee | |
| | SaJJ32 | SA113 | Δ*oatA*::*kan* | | Re-derived from the Δ*oatA::kan* mutant first described by Bera et al | |
| | SaJJ113 | SA113 | *attC*::pJC1111 | | This work | |
| | SaJJ38 | SA113 | Δ*oatA*::*kan*/*attC*::pJC1111 | | This work | |
| | SaJJ39 | SA113 | Δ*oatA*::*kan*/*attC*::pJC1111-*oatA* | | This work | |
| | SaJJ114 | SA113 | Δ*tarM* | | This work | |
| | SaJJ106 | SA113 | Δ*tarS* | | This work | |
| | SaJJ117 | SA113 | Δ*tarMS* | | This work | |
| | SaJJ115 | SA113 | Δ*oatA*::*kan*/Δ*tarM* | | This work | |
| | SaJJ104 | SA113 | Δ*oatA*::*kan*/Δ*tarS* | | This work | |
| | SaJJ118 | SA113 | Δ*oatA*::*kan*/Δ*tarMS* | | This work | |
| | SaJJ123 | SA113 | Δ*oatA*::*kan*/Δ*tarMS* | pOS1-$P_{lgt}$ | This work | |

| | | | | | |
|---|---|---|---|---|---|
| | SaJJ124 | SA113 | Δ*oatA*::*kan*/Δ*tarMS* | pOS1-$P_{lgt}$-*tarM* | This work |
| | SaJJ125 | SA113 | Δ*oatA*::*kan*/Δ*tarMS* | pOS1-$P_{lgt}$-*tarS* | This work |
| | SaJJ126 | SA113 | Δ*oatA*::*kan*/Δ*tarMS* | pOS1-$P_{lgt}$-*tarP* | This work |
| | SaJJ147 | SA113 | Δ*oatA*::*kan*/Δ*tarMS* | pOS1-$P_{lgt}$-*tarM*$_{E403A}$ | This work |
| | SaJJ174 | SA113 | Δ*graRS* | | This work |
| | SaJJ175 | SA113 | Δ*oatA*::*kan*/Δ*graRS* | | This work |
| | SaJJ176 | SA113 | Δ*tarMS*/Δ*graRS* | | This work |
| | SaJJ177 | SA113 | Δ*oatA*::*kan*/Δ*tarMS*/Δ*graRS* | | This work |
| | SaJJ182 | SA113 | Δ*oatA*::*kan* | pOS1-$P_{lgt}$-*sfGFP* | This work |
| | SaJJ3 | JE2 FPR3757 | WT | | BEI |
| | SaJJ11 | JE2 FPR3757 | *agrA*::*bursa* | | BEI; Nebraska Transposon Library |
| | SaJJ21 | JE2 FPR3757 | *agrA*::*bursa*/Δ*oatA*::*kan* | | This work |
| | SaJJ5 | RN4220 | WT | | Originally generated by Richard Novick lab; obtained from Victor J. Torres |
| | SaJJ6 | RN9011 | RN4220/pRN7023 (SaPI integrase, cat194) | | Originally generated by Richard Novick lab (Ruzin et al, 2001); provided by Victor J. Torres |
| | SaJJ35 | LAC | WT | | Victor J. Torres |
| | SaJJ184 | LAC | Δ*oatA* | | This work |

| | | | | | |
|---|---|---|---|---|---|
| | ciJJ11 | ciJJ11 | WT | This work | Isolated from the blood of a patient who developed recurrent MRSA prosthetic valve endocarditis approximately 1.5 weeks after completing a prior course of antibiotic therapy |
| | SaJJ183 | ciJJ11 | Δ*oatA* | This work | |
| | ciJJ26 | ciJJ26 | WT | This work | Isolated from the blood of a patient with extensive vascular grafts who developed recurrent MRSA bacteremia approximately 9 months after completing a prior course of antibiotics for implanted cardioverter-defibrillator infection |
| | SaJJ185 | ciJJ26 | Δ*oatA* | This work | |
| *Staphylococcus carnosus* | SaJJ7 | DSM 20501 | WT | ATCC | |
