## Supplemental Table 2 for "Wall teichoic acid is required for DNA-triggered innate immune receptor activation by *Staphylococcus aureus*"

**Supplemental table 2. Plasmids used in this work**

| Plasmid name | Inserted DNA | Source | Notes |
|---|---|---|---|
| pJC1111 | Empty | Chen et al, Plasmid 2014 | For single-site integration at the *S. aureus* genomic *attC* site |
| pJC1111 | *oatA* with upstream promoter | This work | |
| pJC1111 | *tarM* with upstream promoter | This work | |
| pBursa | N/A | Bae et al, PNAS 2004 | Encodes the mariner-based transposon *bursa aurealis* |
| pMG020 | N/A | Grosser et al, PLoS Pathogens, 2018 | Expression of Himar1 transposase under control of the constitutively active $P_{lgt}$ promoter |
| pIMAY* | Empty | Schuster et al, Microbiology, 2019 | To generate unmarked chromosomal deletions in *S. aureus* |
| pIMAY*-Δ*tarM* | 1kb of each upstream and downstream DNA flanking *tarM* | This work | Required synthesis by Genscript due to complex sequences surrounding tarM |
| pIMAY*-Δ*tarS* | 1kb of each upstream and downstream DNA flanking *tarS* | This work | |
| pIMAY*-Δ*graRS* | 1kb of each upstream and downstream DNA flanking *graRS* | This work | |
| pIMAY*-Δ*oatA* | 1kb of each upstream and downstream DNA flanking *oatA* | This work | |
| pOS1-$P_{lgt}$ | Empty | Bubeck-Wardenburg et al, PNAS, 2006 | Multi-copy plasmid for constitutive expression of genes under control of the $P_{lgt}$ promoter |
| pOS1-$P_{lgt}$-*tarM* | *tarM* | This work | |
| pOS1-$P_{lgt}$-*tarS* | *tarS* | This work | |
| pOS1-$P_{lgt}$-*tarP* | *tarP* | This work | |
| pOS1-$P_{lgt}$-$tarM_{E403A}$ | $tarM_{E403A}$ | This work | |

| | | | |
|---|---|---|---|
| pOS1-sfGFP | Superfolder GFP | This work | sfGFP gene along with its RBS from pSR1037 (Rondthaler et al, ACS Synth Biol 2023) was cloned into pOS1-$P_{lgt}$ |
| pMSCV-IRES-puro | Empty | Our lab | Retroviral vector; puromycin resistance cassette inserted in place of GFP. Generated within our lab |
| pMSCV-IRES-puro-AIM2-mScarlet | AIM2-GGGGS-mScarlet | This work | AIM2 cloned with an upstream RBS and with a C-terminal fusion to a GGGGS flexible linker and mScarlet. |
| pCL-Eco | | Addgene plasmid #12371 | Retroviral packaging plasmid |
| pCMV-VSV-G | | Addgene plasmid #8454 | Pseudotyping plasmid producing vesicular stomatitis virus envelope glycoprotein |
