## Supplemental Table 3 for "Wall teichoic acid is required for DNA-triggered innate immune receptor activation by *Staphylococcus aureus*"

**Supplemental Table 3. Primers used in this work**

| Primer name | Sequence | Notes |
|---|---|---|
| oatA promoter_SalI_F | gactagtcgacGAAAATAACAGTTA | To clone *oatA* with its native promoter into pJC1111 |
| oatA_Stop_KpnI_R | agagtgggtaccTTATTTCTTATTT | To clone *oatA* with its native promoter into pJC1111 |
| tarM_promoter_SalI_F | gactagtcgacaattacacctccaatttcac | To clone *tarM* with its native promoter into pJC1111, excluding the gene upstream of *tarM* |
| tarM_promoter-CDS_R | ccatcataaatattttttcattataattattagcg | To clone *tarM* with its native promoter into pJC1111, excluding the gene upstream of *tarM* |
| tarM_promoter-CDS_F | caattttgttattaagaggtgcgctaataattat | To clone *tarM* with its native promoter into pJC1111, excluding the gene upstream of *tarM* |
| tarM_CDS_SacI_R | AGATGGAGCTCttagctattgaaaag | To clone *tarM* with its native promoter into pJC1111, excluding the gene upstream of *tarM* |
| tarM_pIMAY_5F_NotI | aatgaGCGGCCGCctgtgttactaagtt | To clone ~1kb of each upstream and downstream DNA flanking the *tarM* gene into pIMAY* |
| tarM_pIMAY_5R | cgaaccaaactccgctgctaatacatctcta | To clone ~1kb of each upstream and downstream DNA flanking the *tarM* gene into pIMAY* |
| tarM_pIMAY_3F | ggtatgacctcttcgatgttcaatagagatgta | To clone ~1kb of each upstream and downstream DNA flanking the *tarM* gene into pIMAY* |
| tarM_pIMAY_3R_EcoRI | acatcGAATTCccatcatctgcaatacc | To clone ~1kb of each upstream and downstream DNA flanking the *tarM* gene into pIMAY* |
| tarM_KOcheck_F | gacttcaacactccgtttgattgg | To check for the presence or absence of the *tarM* gene in a putative deletion mutant |
| tarM_KOcheck_R | cgattcatatgccccaatcttatctc | To check for the presence or absence of the *tarM* gene in a putative deletion mutant |
| tarS_pIMAY_5F_NotI | AATGAgcggccgcgaaactttgaataca | To clone ~1kb of each upstream and downstream DNA flanking the *tarS* gene into pIMAY* |
| tarS_pIMAY_5R | cacaatgatttgagggcatttattatttaaatttc | To clone ~1kb of each upstream and downstream DNA flanking the *tarS* gene into pIMAY* |
| tarS_pIMAY_3F | gtcaaagtgggagaggtataatgatgaaatt | To clone ~1kb of each upstream and downstream DNA flanking the *tarS* gene into pIMAY* |
| tarS_pIMAY_3R_EcoRI | AGATGgaattccatgtaaaattcatgaata | To clone ~1kb of each upstream and downstream DNA flanking the *tarS* gene into pIMAY* |
| tarS_KOcheck_F | catcagactctagaccaacgatgtc | To check for the presence or absence of the *tarS* gene in a putative deletion mutant |
| tarS_KOcheck_R | gatagcacgatagcacctcagtg | To check for the presence or absence of the *tarS* gene in a putative deletion mutant |
| pIMAY_oatA_5F_NotI | aatgaGCGGCCGCggtCTTATAA | To clone ~1kb of each upstream and downstream DNA flanking the *oatA* gene into pIMAY* |
| oatA_pIMAY_5'_R | CAAAAGTTTAGTGCATCAAATT | To clone ~1kb of each upstream and downstream DNA flanking the *oatA* gene into pIMAY* |
| oatA_pIMAY_3'_F | GAGAAAAATAAATGGGGCGTT | To clone ~1kb of each upstream and downstream DNA flanking the *oatA* gene into pIMAY* |
| pIMAY_oatA_3R_SalI | agagtgGTCGACgcaACATGACC | To clone ~1kb of each upstream and downstream DNA flanking the *oatA* gene into pIMAY* |
| oatA_KOcheck_F | ctgagtacactagatacaaactaaacg | To check for the presence or absence of the *oatA* gene in a putative deletion mutant |
| oatA_KOcheck_R | gtccatagttcatttctgaaacacg | To check for the presence or absence of the *oatA* gene in a putative deletion mutant |
| pOS1_tarM_RBS_BamHI_F | agatcGGATCCgaggtgaacccAtga | To clone *tarM* into pOS1-$P_{lgt}$ |
| pOS1_tarM_PstI_R | AGATGctgcagTtagctattgaaaagatt | To clone *tarM* into pOS1-$P_{lgt}$ |
| pOS1_tarS_RBS_BamHI_F | agatcGGATCCgaggtgaacccAtga | To clone *tarS* into pOS1-$P_{lgt}$ |
| pOS1_tarS_SphI_R | AGATGgcatgcTtatttatttagtggataa | To clone *tarS* into pOS1-$P_{lgt}$ |
| pOS1_tarP_RBS_BamHI_F | agatcGGATCCgaggtgaacccATG | To clone *tarP* into pOS1-$P_{lgt}$ |
| pOS1_tarP_PstI_R | AGATGctgcagCTATAATAGCTT | To clone *tarP* into pOS1-$P_{lgt}$ |
| tarM_E403A_R | catacttaacccttgacctgcatattgagatg | To generate a catalytically inactive but structurally intact E403A mutant of *tarM* |
| tarM_E403A_F | cgtttctacatctcaatatgcaggtcaagggt | To generate a catalytically inactive but structurally intact E403A mutant of *tarM* |
| AIM2-mScarlet gBlock | gagtagatctGATCCTGGGACTGT | To clone an AIM2-mScarlet fusion protein into the retroviral vector pMSCV-IRES-puro |
| SA_qPCR_16S_F | ctcgtgtcgtgagatgttgg | For qPCR of *S. aureus* 16S |
| SA_qPCR_16S_R | GTTTGTCACCGGCAGTCAAC | For qPCR of *S. aureus* 16S |
| qPCR_tarO_F | gtcaaattgccgctgccttag | For qPCR of *S. aureus* *tarO* |
| qPCR_tarO_R | GCCAAACCATCGAGTCCATC | For qPCR of *S. aureus* *tarO* |
| SA_qPCR_tarM_F | GGCCTGTTGTTGCCTTTGAC | For qPCR of *S. aureus* *tarM* |
| SA_qPCR_tarM_R | CGCTTTCGAACCAAACTCCG | For qPCR of *S. aureus* *tarM* |
| qPCR_tarS_F | ctcgtacgaatggcttctcac | For qPCR of *S. aureus* *tarS* |
| qPCR_tarS_R | CTCGCGTAGTGCAACAATGG | For qPCR of S. aureus *tarS* |
| mm_ActinIntron_F | TGG CCT TGG AGT GTG TAT | For qPCR of mouse β-actin |
| mm_ActinIntron_R | AGG GCA GGT GAA ACT GTA | For qPCR of mouse β-actin |
