## Supplemental Table 4 for "Wall teichoic acid is required for DNA-triggered innate immune receptor activation by *Staphylococcus aureus*"

**Supplemental table 4. CRISPR sgRNAs used in this work**

| sgRNA name | Sense/Antisense | PAM | Target sequence | Source |
|---|---|---|---|---|
| mLYZ1.1.AC | - | TGG | AATGGAATGGATGGCTACCG | IDT |
| mLYZ2.1.AA | + | TGG | CTCACAACGTTCATAGACCT | IDT |
| mAIM2.1.AB | + | AGG | CGGCCTGGACCACATCACGG | IDT |
